# Gene Supplementation of *MYO7A* or activation of *Myo7b* for treatment of Usher syndrome 1B

**DOI:** 10.64898/2026.07.02.736025

**Authors:** David M. Mittas, Dina Y. Otify, Zoran Gavrilov, Thomas Heigl, Jan Suchomski, Paulina Deltuvaite, Klara Hinrichsmeyer, Olivier Mercey, Franz Kynast, Jan Motlik, Zdenka Ellederova, Taras Ardan, Andreas Klingl, Jennifer Grünert, Verena Mehlfeld, Anastasiia Kolesnikova, Ruslan Nyshchuk, Jana Juhasova, Stefan Juhas, Saskia Drutovic, M. Dominik Fischer, Miroslav Veith, Zbynek Stranak, Nanda Boon, Jan Wijnholds, Andreas Wiest, Pavel Kielkowski, Gülce Gökce, Paul Guichard, Virgine Hamel, Hermann Ammer, Stylianos Michalakis, Susanne Koch, Martin Biel, Elvir Becirovic

## Abstract

Mutations in *MYO7A* result in the most severe subtype of Usher syndrome, the leading genetic cause of deafblindness. The large size of *MYO7A* requires dual adeno-associated virus (AAV) vectors for gene transfer or alternative methods to treat retinal defects. Here, we evaluated two treatment approaches: i) Supplementation of the human *MYO7A* gene via dual mRNA trans-splicing AAVs, and ii) CRISPR/Cas-mediated activation of the related murine *Myo7b* gene. Upon *MYO7A* supplementation, the transgenic MYO7A transcript and protein were expressed and correctly localized in retinal pigment epithelial (RPE) and photoreceptors of mice, pigs, and human retinal organoids. In RPE-and photoreceptor-specific *Myo7a* knockout mice, we could restore MYO7A expression and localization of melanosomes in RPE cells to wild-type levels. *Myo7b* activation led to partial restoration of melanosome localization, and the localization of MYO7B protein was largely comparable to MYO7A. These findings indicate that both approaches are in principle suitable for the therapy of Usher syndrome.

## Introduction

With a prevalence of approximately 1:6,000-10,000, Usher syndrome (USH) is the most common cause of genetic deafblindness worldwide^1–3^. Depending on the severity of the disease, three subtypes are distinguished (USH1-3). USH1 is clinically the most severe subtype and is characterized by congenital deafness, vestibular dysfunction, and usually early-onset, progressive retinal defects due to retinitis pigmentosa^4–7^. Of the six USH1 genes identified to date, mutations in *MYO7A* (USH1B) are by far the most common cause of this subtype, accounting for >50% of cases^8–12^. *MYO7A* encodes Myosin 7a, an unconventional motor protein found primarily in the ciliary and periciliary membranes of photoreceptors, in the apical region of retinal pigment epithelial (RPE) cells, and in the stereocilia of sensory hair cells^13–15^. Myosin 7a is responsible for the transport of melanosomes in RPE cells and for the structural integrity of hair cells in the cochlea and photoreceptors^16–21^. Myosin 7a is embedded in a network of various proteins through direct or indirect protein-protein interactions, many of which are associated with USH^1,5,22,23^.

While hearing defects can be treated relatively well with cochlear implants, there are no effective therapies for retinal defects. The coding sequence of *MYO7A* (approx. 6.7 kb) exceeds the DNA capacity of adeno-associated virus (AAV) vectors, the currently most used platform for *in vivo* gene therapy. Strategies that can overcome this size limitation, such as dual AAVs are therefore necessary for the delivery of *MYO7A* into the target retinal cells^24^. Apart from gene supplementation, CRISPR/Cas-mediated activation (CRISPRa) of related genes that have the potential to take over the function of the defective gene represents another alternative for the treatment of genetic diseases^25–30^. With its high similarity to Myosin 7a at the structural and sequence level, Myosin 7b (MYO7B) fulfills this criterion^31^. Moreover, MYO7B has been shown to interact in the microvilli of intestines with highly similar adaptor proteins as MYO7A in stereocilia and forms a comparable protein network complex to the Usher-network^32–35^.

To investigate MYO7A function and test potential therapies, the naturally occurring Shaker mouse model for USH1B has been used to date. This mouse model exhibits a weak retinal and a strong cochlear and vestibular phenotype, which is characterized by typical circling behavior^36–39^. The most established ocular phenotype in this mouse is the mislocalization of melanosomes in the RPE cells and is therefore widely used to test the efficacy of retinal gene therapies^40,41^.

In this study, we developed and evaluated the supplementation of human *MYO7A* or the activation of the endogenous murine *Myo7b* gene in a conditional USH1B mouse model that shows a similar ocular phenotype as the Shaker mouse.

## Results

### Development of a MYO7A gene supplementation therapy using REVeRT

To supplement human *MYO7A*, we used dual mRNA trans-splicing (REVeRT) AAV vectors from our recently published study^24,31^. To this end, we first directly compared three split sites within the coding *MYO7A* sequence regarding the reconstitution efficiency of the *MYO7A* transcript in HEK293 cells co-transfected with the AAV plasmids (Extended Data Fig. 1a-c). The split site with the highest reconstitution efficiency (b) was used for subsequent experiments. We further transfected 661W cells as retinal cell line and MEF cells, which do not endogenously express *MYO7A*, to confirm correct splicing between the two *MYO7A* halves (Extended Data Fig. 1d). Subsequently, the reconstitution efficiency of the dual REVeRT vectors was evaluated *in vivo* in the retina of wild-type mice and minipigs as well as in wild-type human retinal organoids. As MYO7A is expressed in both RPE and photoreceptors, it is plausible to assume that both cell types would need to be targeted for effective therapy in USH1B patients. Since there is currently no therapeutically applicable promoter that is compatible with AAV vectors and specifically expressed in these two cell types, we used two common, ubiquitous promoters for the initial experiments in mice and retinal organoids: CMV (cytomegalovirus) and CAG (CMV enhancer fused to the chicken beta-actin promoter). Unless otherwise stated, we used the AAV8_Y733F capsid for all subsequent experiments^42^. After subretinal injection of wild-type mice with dual REVeRT AAV vectors (Fig. 1a), we observed 68% reconstitution efficiency (Fig. 1b). To address the MYO7A protein expression, a myc-tag and the corresponding antibody were used to distinguish between the transgenic and the endogenous MYO7A protein. With this antibody, we only detected a specific signal at the size of the full-length MYO7A protein (approx. 250 kDa) in eyes that had been co-injected with dual REVeRT AAV vectors (Fig. 1c). To quantify the protein expression of transgenic MYO7A in injected eyes, semi-quantitative western blot analysis was performed using an antibody that recognises the N-terminus of both the mouse and human MYO7A. This resulted in 56% ± 3.54% expression of transgenic MYO7A relative to the endogenous protein in wild-type mice (Fig. 1d). Moreover, we detected strong expression of transgenic MYO7A in the photoreceptors and RPE of mice at the protein and transcript level for both promoters (Fig. 1e-f). The localization of transgenic MYO7A was mostly detectable in photoreceptors and RPE cells (Fig. 1g). For transduction of wild-type human retinal organoids, we used the previously described AAV2.NN capsid^43^ (Fig. 2a). Efficient reconstitution of transgenic MYO7A was observed at the transcript and protein levels (Fig. 2b-d). The expression was largely restricted to the photoreceptors (Fig. 2e). In injected retinas of minipigs, the full-length transgenic MYO7A was detectable (Fig 2f-g), leading to 30% transgenic MYO7A expression relative to the endogenous protein (Fig. 2h-i). Like in mice, a comparably strong signal for transgenic MYO7A was also detectable in the inner nuclear layer in addition to the photoreceptors and RPE cells (Fig. 2j). Another cohort of pigs was subretinally injected with dual AAVs using the previously described AAV2.GL capsid^43^, expressing MYO7A under the control of the CAG promoter (Fig. 2k). In this experiment, we tested two different AAV vector doses (5×10^11^ or 1×10^12^ total viral genomes (vg)). Six weeks post-injection, we detected reconstituted transgenic MYO7A at both the transcript and protein levels with a slightly increased expression in the higher dose (Fig. 2l-n).

**Fig. 1.**
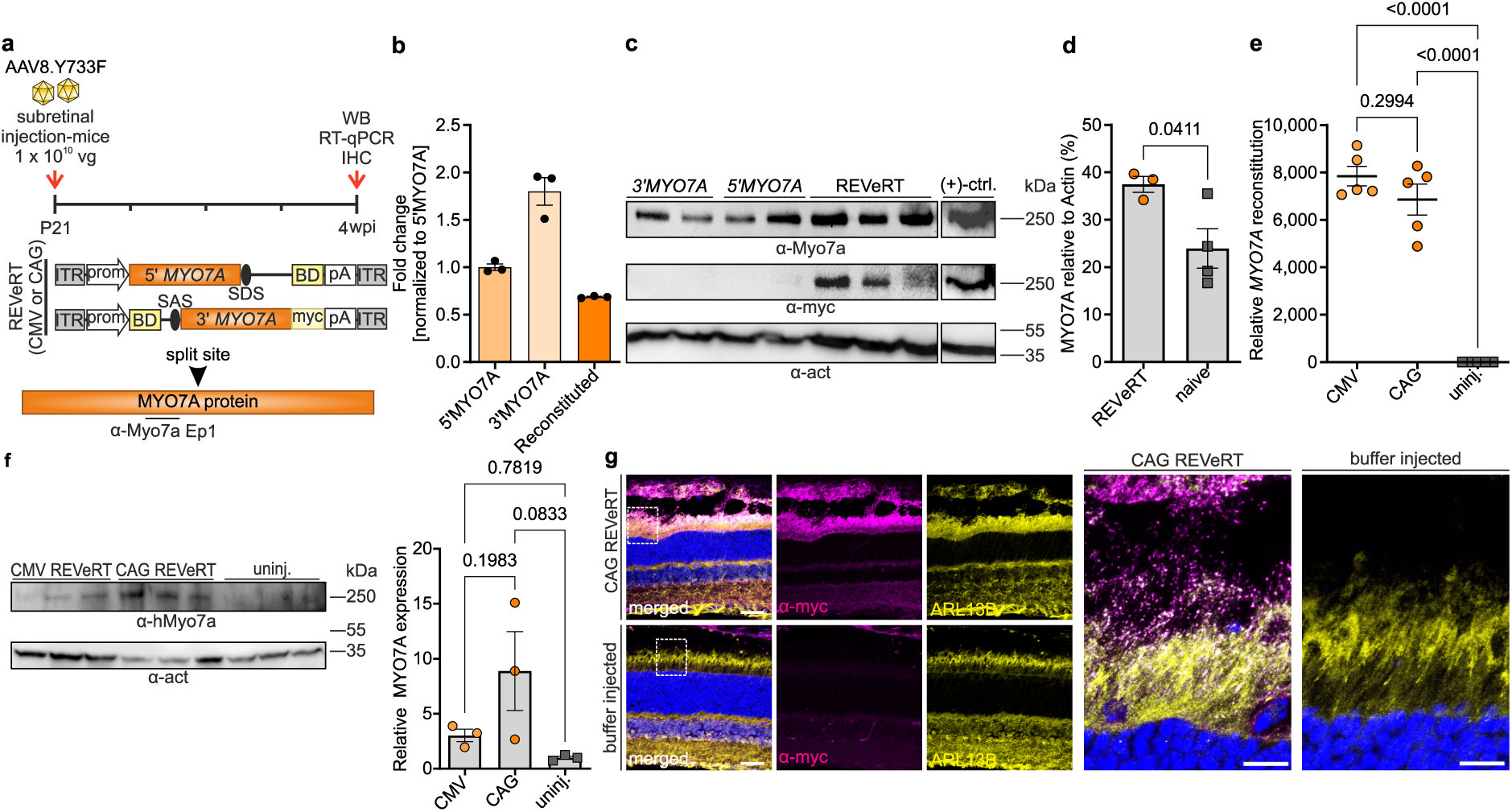
Validation of *MYO7A* supplementation in wild-type mice. *a,* Experimental design depicting the timeline for subretinal injections of C57BL/6J mice and the expression cassettes used. P21, postnatal day 21. wpi, weeks post-injection. Ep1, Schematic depiction of the binding site of the α-Myo7a antibody epitope. WB, western blotting. IHC, immunohistochemistry. ITR, inverted terminal repeats. BD, binding domain. pA, polyadenylation signal. SDS, SAS, splice donor or acceptor sites. *b,* Quantitative PCR analysis 4 weeks post-injection showing reconstitution efficiency, normalized to the relative abundance of the 5′ *MYO7A* transcript (n = 3; data adapted from Mittas et al. 2026^24^). *c,* Western blot using whole eyecup lysates 4 weeks after injection, probed with anti-myc (α-myc) and a MYO7A antibody (α-Myo7a) recognizing both human and mouse protein (hereafter referred to as pan-M7A). Eyes injected with single AAV halves served as negative controls (n = 3 for REVeRT & n = 4 for the control), and HEK293T lysates transfected with full-length myc-tagged MYO7A as positive control. *d,* Semi-quantitative analysis of c; MYO7A levels were normalized to β-actin (α-act). *e,* Relative MYO7A expression in whole eyecups (n = 5) after injection of dual AAVs in presence of the CAG or CMV promoter, determined by qPCR using split site spanning primers. Uninjected eyes (n = 5) served as negative controls. *f,* Western blot of whole eyecup lysates using a human-specific MYO7A antibody (α-hMyo7a, n = 3). Right panel shows the semi-quantitative analysis; MYO7A levels were normalized to β-actin. *g,* Representative immunostaining of retinas (n = 2 per group) injected with dual AAVs or with formulation buffer; Right panels, Close-up images of the regions marked by dashed rectangles. ARL13B marks primary cilia. Scale bar, 30 μm. Statistical analysis was performed using a two-tailed unpaired t test with Welch’s correction (*d*) and one-way ANOVA followed by Tukey’s post hoc test (*e, f*). Scatter plots show mean ± s.e.m. All source data are provided in the Source Data file.

**Fig. 2.**
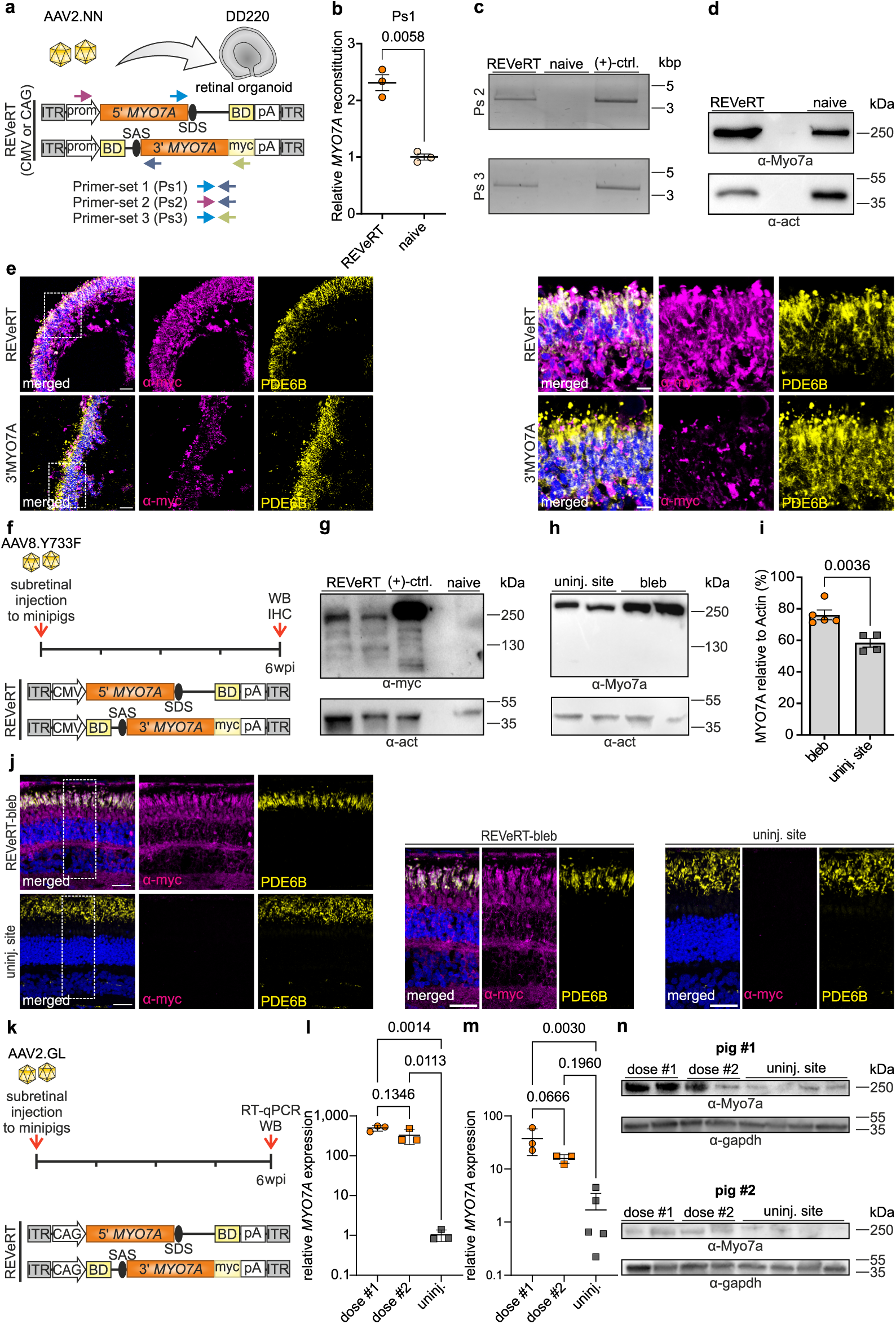
***MYO7A* supplementation in wild-type human retinal organoids and pigs.** *a,* Experimental workflow. Human retinal organoids were transduced at differentiation day (DD) 220 with dual AAVs expressing one half of *MYO7A* using the AAV2.NN capsid as indicated. The binding positions of the individual primer sets (ps1-3) in the expression cassettes used for experiments in b and c are indicated by colored arrows. *b,* Relative *MYO7A* expression in transduced versus naïve organoids measured by qPCR with split site spanning primers (n = 6 organoids per group; two organoids pooled per datapoint). *c,* RT–PCR amplification of reconstituted transcripts using another two split site spanning primer pairs. cDNA from HEK293T cells transfected with myc-tagged full-length MYO7A served as positive control. *d,* Representative western blot of organoid lysates probed with a pan-M7A antibody. *e,* Representative immunostaining of organoids transduced with dual AAV vectors expressing both MYO7A halves or with AAVs expressing 3′MYO7A alone. Right panels, Close-up images. Scale bar, 30 µm. *f,* Schematic overview of experiments in 3 months old wild-type minipigs (n = 2). Subretinal injection was performed with AAV8Y733F capsids (1 × 10¹² total vg). *g,* Western blot of retinal punches from the bleb region probed with an anti-myc antibody. HEK293T lysates transfected with a full-length myc-tagged MYO7A expression cassette served as positive control. *h,* Representative western blot of punches from the bleb and uninjected regions probed with a pan-M7A antibody. *i,* Semi-quantitative analysis of western blot of punches from the bleb and uninjected regions probed with pan-M7A antibody; each datapoint represents one individual retinal punch from two pigs. *j,* Representative immunostaining of the subretinal bleb and uninjected regions probed with an anti-myc antibody. Right panels show close-up images. Scale bar, 30 µm. *k,* Schematic overview of experiments in another cohort of 3 months old wild-type minipigs (n = 2). Pigs were subretinally injected with dual AAVs using the AAV2.GL capsid. One eye received 1×10^12^ total vg (dose #1), while the contralateral eye received 5×10^11^ total vg (dose #2). *l,m,* Relative *MYO7A* expression (l, pig #1, m, pig #2) in the injected bleb area versus the uninjected site determined by RT-qPCR with split site spanning primers. Each datapoint represents a biopsy punch from the respective injection region. *n*, Western blot of retinal punches from the bleb region and the uninjected site probed with a pan-M7A antibody (upper panel, pig #1, lower panel, pig #2). Statistical analysis was performed using a two-tailed unpaired t test with Welch’s correction (*b, i*) or one-way ANOVA followed by Tukey’s post hoc test (*l, m*). Scatter plots show mean ± s.e.m. All source data are provided in the Source Data file.

### Side-by-side comparison of REVeRT MYO7A and DNA trans-splicing vectors

With dual REVeRT AAV vectors, the split gene is mainly reconstituted at the mRNA level with the help of mRNA trans-splicing^24,31^. In previous studies in Shaker mice and in the recently initiated clinical trial in USH1B patients, dual AAV vectors for *MYO7A* supplementation were used in which the reconstitution of *MYO7A* occurs at the genomic level (hereinafter referred to as DNA trans-splicing)^40,41,44–46^. We compared the performance of dual REVeRT AAVs directly with their DNA trans-splicing counterparts and observed approximately threefold higher reconstitution efficiency of REVeRT at the transcript and protein levels (Extended Data Fig. 2a-d). In a preliminary dose-finding expression study, we detected a strong MYO7A signal at the protein levels for three different AAV doses (5×10^9^, 1×10^10^, and 5×10^10^ total vg) in eyes of subretinally injected wild-type mice (Extended Data Fig. 2e). The transgenic full-length MYO7A protein could be detected for all individual doses in a dose-dependent manner. However, the high dose did not show higher MYO7A expression compared to the mid dose, (Extended Data Fig. 2f-g), which is why the latter was defined as the therapeutic dose for subsequent experiments in the USH1B mouse model.

### Establishment and characterization of a novel conditional Myo7a-KO-mouse model

Next, we set out to provide proof of concept for the *MYO7A* supplementation therapy using dual REVeRT AAV vectors in the USH1B mouse model. We decided not to use the Shaker mouse model, as it exhibits severe behavioral abnormalities due to the vestibular defects and therefore represents a high level of stress under animal welfare legislation. To create an RPE and retina specific *Myo7a* knockout, we took advantage of the recently published conditional USH1B mouse model in which exons 10-11 are flanked by loxP sites. leading to a deletion of 197 bp in the coding sequence in presence of Cre, followed by a premature stop codon^47^. We crossbred this line with the Cre reporter line under the control of the Rx (Retina and Anteria Neural Fold Homebox) promoter (Extended Data Fig. 3a), which is expressed in precursors of photoreceptor and RPE cells^48,49^. In the resulting Rx-Cre(+)-Myo7a (hereafter referred to as Cre(+)) line, we analyzed the expression of MYO7A at the transcript and protein levels. We separated the neuronal retina from the RPE and analyzed both tissues individually. Using primers targeting the deleted genomic region in the *Myo7a* locus, we observed virtually complete recombination in the retina at both the genomic and transcript levels (Extended Data Fig. 3b-c). However, although the deletion was predicted to cause a frameshift and an early stop codon in the *Myo7a* transcript (Extended Data Fig. 3c), residual MYO7A protein was still detectable, albeit at lower levels compared to the Cre(-) controls (Suppl. Fig 3d-e). By comparison, RPE samples showed a partial knockout at both transcript and protein levels, indicating mosaicism (Extended Data Fig. 3f-i).

In subsequent quantitative proteomics we could detect MYO7A peptides in both the RPE and retinal samples. As expected, the levels of MYO7A peptides were reduced in Cre(+) animals in both cell types. Moreover, in line with previous findings, the overall MYO7A protein expression was substantially higher in RPE cells (Extended Data Fig. 3j). In RPE cells, we identified a peptide from the deleted region of MYO7A, which was reduced to a higher extent than total MYO7A protein levels (Extended Data Fig. 3j-k). In a separate experiment we isolated the lysates from the whole eyecups containing both retina and RPE cells and observed intermediate reductions in MYO7A transcript and protein levels (Extended Data Fig. 3l-o). In addition, we performed western blot analysis of protein lysates that had been digested by endogenous proteases, as described in our recent study^50^. Using two MYO7A antibodies recognizing different epitopes, we identified a different band pattern for Cre(+) RPEs compared to their Cre(-) counterparts (Extended Data Fig. 3p). This indicates that at least some of the residual MYO7A expression in Cre(+) retinas might be due to the expression of a truncated protein.

In immunostainings, a clustered knockout of MYO7A was visible in the RPE cells (Extended Data Fig. 4a), which is most likely due to mosaicism in these cells, as described in previous studies on other mouse models^48,49^. To analyze the subcellular localization of MYO7A within photoreceptor cells more precisely, we employed expansion microscopy^51,52^. In Cre(-) animals, the MYO7A signal was detected at the ciliary rootlet in the inner segment. Additional MYO7A was found to surround the basal body of the mother and daughter centriole, and in the transition fibers of the photoreceptor centriole (Extended Data Fig. 4b-c). In comparison, the MYO7A signal in Cre(+) animals was completely abolished at the rootlet and significantly reduced in the basal body and transition fiber structures (Extended Data Fig. 4d-e). The decrease was consistent with the western blot results from the retinal lysates described above (cf. Extended Data Fig. 3e). Electroretinography and OCT measurements at 5 and 12 months of age showed no differences between Cre(-) and Cre(+) animals (Extended Data Fig. 4f-j). This is in line with the previously published results from the Shaker or other USH mouse models^36,47^.

In Cre(+) animals, we observed a significant mislocalization of melanosomes in apical parts of RPE cells, which is similar to the observations from the Shaker mouse (Extended Data Fig. 5a-c). This mislocalization occurred in clusters, consistent with the RPE mosaicism described above. There is currently conflicting data on whether MYO7A knockout leads to defective transport of opsins^14,15,21,53^. Under the conditions tested herein, we found no evidence of defective transport of opsins in Cre(+) mice (Extended Data Fig. 5d). Based on these results, we used melanosome localization in RPE cells and the subcellular localization of MYO7A within the ciliary region of photoreceptors as two parameters to evaluate the success of gene supplementation therapy in Cre(+) mice.

### Gene therapy of Cre(+) mice with MYO7A supplementation

For therapy, we injected the animals with dual REVeRT AAVs expressing *MYO7A* under the control of the CAG promoter on postnatal day 21 (P21). The contralateral eye of each mouse served as a control and remained uninjected (Fig. 3a). Four (Fig. 3a-g) and 20 weeks (Fig. 4b-d) after injection, we detected an increase in MYO7A expression at the transcript and protein levels of transduced retinas and RPEs compared to control samples. The total expression of MYO7A was restored to the level of Cre(-) mice in both RPE cells and the retina (Fig. 3b-g & Fig. 4e-f). In quantitative proteomics analysis of Cre(+) RPEs we identified an increase in photoreceptor outer segment proteins compared to Cre(-) animals (Fig. 3h,l). This indicates a defective transport of RPE phagosomes, which incorporate photoreceptor outer segments, from the apical to the basal RPE for lysosomal degradation as described previously for Shaker mice^21,54–57^. Subretinal injection of the dual AAVs expressing *MYO7A* led to a complete restoration of outer segment protein expression in these mice (Fig. 3h-l).

**Fig. 3.**
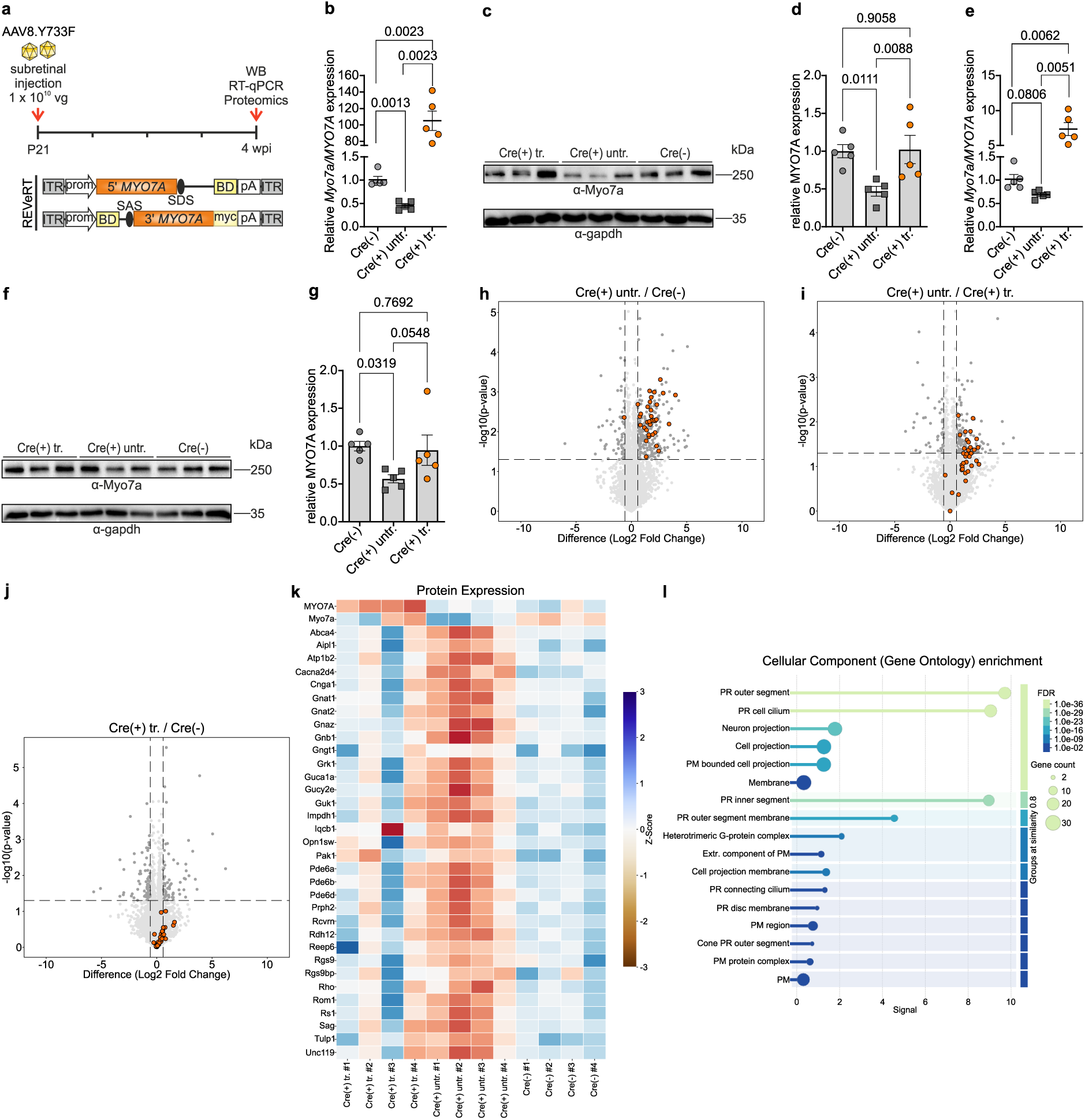
Analysis of protein and gene expression upon *MYO7A* gene supplementation in Cre(+) mice. *a,* Experimental overview. Cre(+) mice were subretinally injected with dual AAVs in one eye, the contralateral eye remained uninjected. Age-matched Cre(–) littermates served as controls. P21, postnatal day 21. *b,* Relative *Myo7a* expression in the neuronal retina of treated, untreated, and Cre(–) control animals measured by qPCR with split site spanning primers targeting mouse and human *Myo7a/MYO7A* transcripts (n = 5 per group). *c,* Representative western blot of retinal lysates from treated, untreated, and Cre(–) animals. *d,* Semi-quantitative analysis of retinal MYO7A protein expression normalized to GAPDH (n = 5 per group). *e,* Relative *Myo7a/MYO7A* expression in RPE of treated, untreated, and Cre(–) control animals (n = 5 per group). *f,* Representative western blot of RPE lysates. *g,* Semi-quantitative analysis of RPE MYO7A protein expression normalized to GAPDH (n = 5 per group). *h-j,* Volcano plots showing pairwise comparisons between i) untreated Cre(+) and Cre(-) RPEs (h) ii) untreated Cre(+) and Cre (+) treated with *MYO7A* supplementation, and iii) treated Cre(+) and naïve Cre(-) RPEs, where photoreceptor outer segment proteins are labeled. *k*, Heatmap showing the protein expression of the differentially regulated protein clusters in RPE (labeled h-j). Differentially expressed proteins were defined by |logFC| ≥ 0.58 and an adjusted p-value < 0.05 (cut = 1.3). Upregulated photoreceptor proteins in RPEs of untreated Cre(+) eyes (*h,i*) are highlighted in orange. *l*, Diagram showing the cellular component (Gene Ontology) enrichment of the identified upregulated cluster of proteins in untreated Cre(+) RPEs (n = 4 per group). PM, photoreceptor membrane. Extr, extracellular. Statistical analysis: one-way ANOVA with Tukey’s post hoc test (*b, d, e, g*). Scatter plots show mean ± s.e.m. All source data are provided in the Source Data file.

**Fig. 4.**
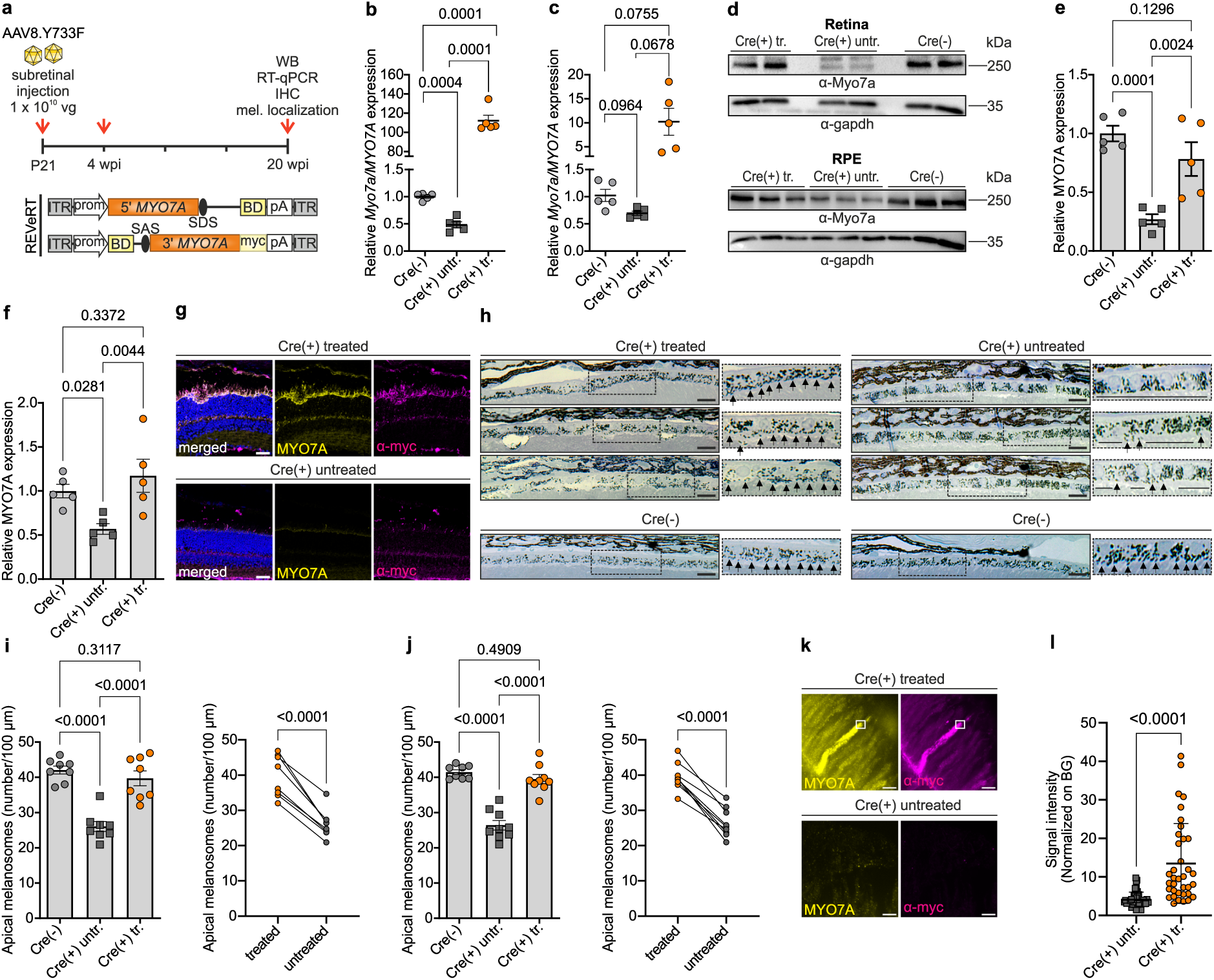
***MYO7A* gene supplementation therapy in Cre(+) mice.** *a,* Experimental overview. Cre(+) mice were subretinally injected with dual AAVs in one eye, the contralateral eye remained uninjected. Age-matched Cre(–) littermates served as controls. *b, c,* Relative *Myo7a* expression in neuronal retina (*b*) and RPE (*c*) of treated, untreated, and Cre(–) control animals measured by qPCR with split site spanning primers binding both mouse *Myo7a* and human *MYO7A* transcripts (n = 5 per group). *d,* Representative western blot of retinal and RPE lysates from treated, untreated, and Cre(–) control animals. *e, f,* Semi-quantitative analysis of MYO7A expression in the retina (*e*) and RPE (*f*), normalized to GAPDH (n = 5 per group). *g,* Representative immunostainings of Cre(+) mice 5 months after injection of dual AAVs displayed in a, showing injected versus uninjected regions. Scale bar, 30 µm. *h,* Representative semi-thin (500 nm) retinal cross-sections from treated or untreated Cre(+) and Cre(–) mice, stained with epoxy-tissue stain. Arrows highlight melanosomes in apical processes; horizontal lines indicate areas lacking apical melanosomes. Scale bar, 20 µm. *i, j,* Double-blinded quantification of apical melanosomes across the RPE surface, normalized to 100 µm RPE length, 4 weeks (*i*) (n = 8 per group) and 20 weeks (*j*) (n = 9 for Cre(+) and 8 for Cre(-)) post-injection. Right panels show paired comparisons of treated versus contralateral untreated eyes. *k,* Representative immunostainings of injected and expanded tissues. Scale bar, 1 µm. *l,* Quantification of MYO7A signal intensity at the ciliary basal body, normalized to background. Statistical analysis: two-tailed paired t test (*i, j*, right panels); two-tailed unpaired t test with Welch’s correction (*l*); and one-way ANOVA with Tukey’s post hoc test (*b, c, e, f,* and left panels of *i, j*). Scatter plots show mean ± s.e.m. All source data are provided in the Source Data file.

The MYO7A protein was detectable in immunostainings of retinal cryosections in both RPE cells and photoreceptors of all treated eyes (n = 3) (Fig. 4g & Extended Data Fig. 6a). Four and 20 weeks after injection treated Cre(+) retinas exhibited virtually no mislocalization of melanosomes in the injected RPE area (Fig. 4h). In fact, the melanosome localization in the treated eyes was statistically indistinguishable from that of the Cre(-) control animals (Fig. 4i-j). Compared to the Cre(-) mice, in the expansion microscopy of treated Cre(+) retinas, a much stronger MYO7A signal was detected in the entire inner segment and at the upper base of the inner segments corresponding to the basal body and transition zone of photoreceptors. Due to this strong expression of transgenic MYO7A, no distinction between the basal body, rootlet, and the transition zone was possible (Fig. 4k). MYO7A signal within the entire region of the upper inner segment, basal body, and transition fibers showed an increase compared to untreated contralateral eyes of Cre(+) animals (Fig. 4k-l & Extended Data Fig. 6b-c).

Subretinal or intravitreal injections are currently the main delivery routes for therapy of retinal diseases, both with specific advantages and disadvantages^58–60^. To test whether *MYO7A* can be reconstituted in the mouse retina after intravitreal injection, we administered the recently published AAV2.GL vectors in combination with our dual REVeRT AAVs (Extended Data Fig. 7a). Compared to subretinal injection, intravitreal application resulted in poorer reconstitution efficiencies and less consistent results but still showed a clearly detectable MYO7A signal at the transcript and protein levels (Extended Data Fig. 7b-d).

In summary, subretinal delivery of our dual mRNA trans-splicing AAVs to a novel conditional mouse USH1B model leads to high levels of MYO7A protein and transcript expression and restores the key molecular phenotype in RPE cells.

### Gene therapy of Cre(+) mice by transcriptional activation of Myo7b

In the second part of the study, we evaluated the efficiency of CRISPRa-mediated activation of the murine *Myo7b* gene in Cre(+) mice. We identified MYO7B as a potential functional counterpart of MYO7A, based on strong conservation of functional domain architecture and emerging evidence of shared interaction partners (Extended Data Fig. 8a-e). To this end, we used the recently published *Myo7b* sgRNAs in combination with dead(d)Cas9-VPR, which enabled efficient *Myo7b* activation *in vitro* and *in vivo*^31^.

For therapy, we subretinally injected Cre(+) animals on P21 with dual REVeRT AAVs expressing the 5’ and 3’ parts of the dCas9-VPR coding sequence under the control of the CAG promoter (Fig. 5a). The contralateral eye of each mouse served as a control. Four and 20 weeks after injection, we detected an increase in *Myo7b* expression at the transcript and protein levels compared to uninjected eyes (Fig. 5b-g). In line with that, we detected reconstituted dCas9-VPR at the transcript and protein levels exclusively in injected eyes (Fig. 5d-g & Extended Data Fig. 9a-d). The MYO7B protein signal was mainly observed in the inner segments of photoreceptors and in RPE cells (Fig. 5h). It was not possible to determine the exact subcellular localization of MYO7B in the inner segments of the photoreceptors, as none of the antibodies tested worked in expansion microscopy (data not shown).

**Fig. 5.**
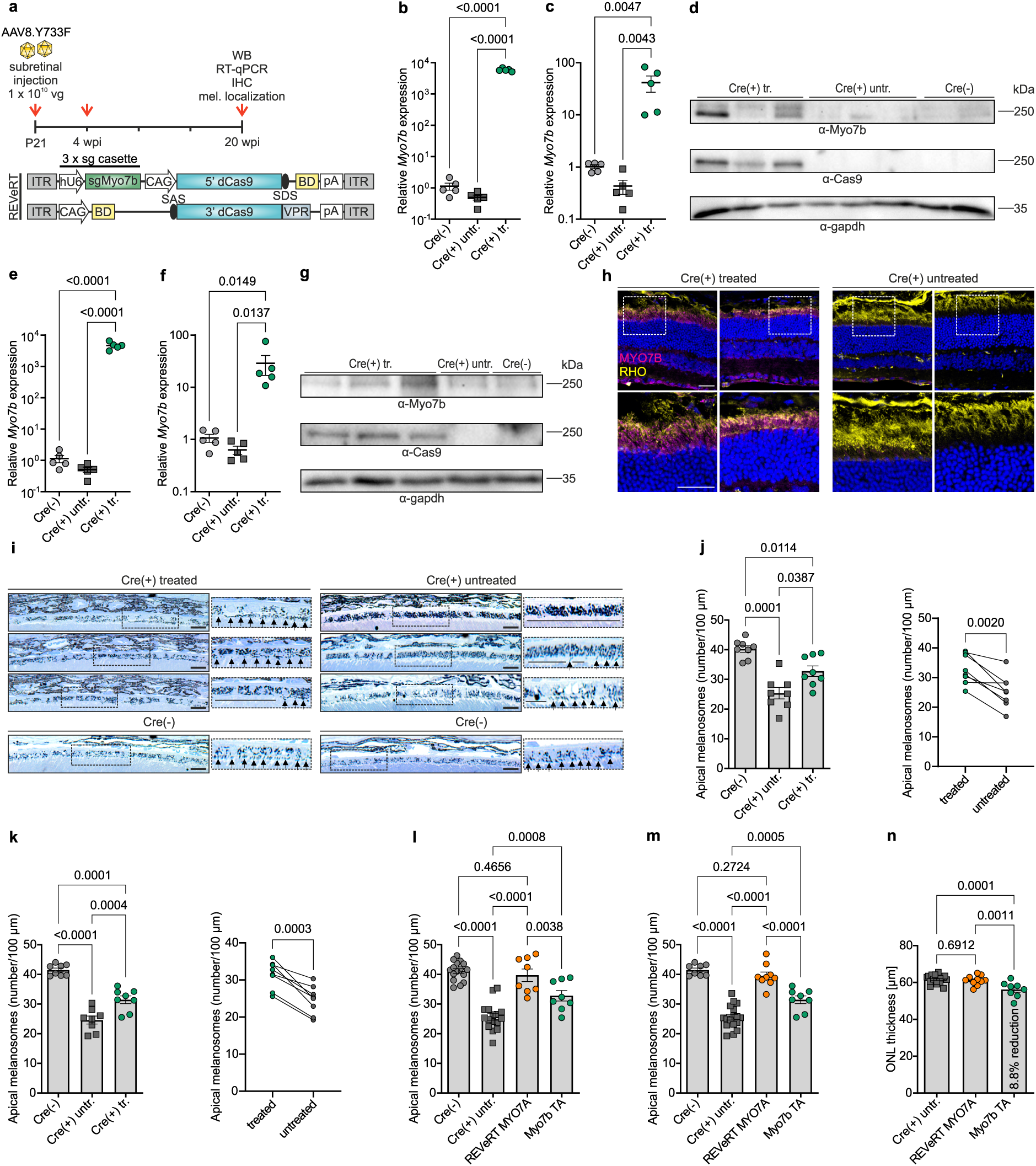
***Myo7b* activation in Cre(+) mice.** *a,* Experimental overview. One eye of Cre(+) mice was subretinally injected with dual AAV vectors expressing the split dCas9–VPR and the triple sgRNA cassette targeting the promoter of the murine *Myo7b* gene; The contralateral eyes remained non-injected and served as paired controls. Age-matched Cre(–) littermates served as another control group. *b, c,* Relative *Myo7b* expression in neuronal retina (*b*) and RPE (*c*) 4 weeks post-injection (n = 5 per group). *d,* Representative western blot of treated or untreated, Cre(+) and Cre(–) retinas 4 weeks post-injection. *e, f,* Relative *Myo7b* expression in neuronal retina (*e*) and RPE (*f*) 20 weeks post-injection (n = 5 per group). *g,* Representative western blot of treated or untreated Cre(+), and untreated Cre(–) retinas 20 weeks post-injection. *h,* Representative immunostaining of another cohort of Cre(+) mice (n = 2) injected with dual AAV vectors displayed in a, showing injected and uninjected regions 20 weeks post-injection. Lower panels show the corresponding close-up images. Rhodopsin served as a marker for rod outer segments. Scale bar, 30 µm. *i,* Representative semi-thin (500 nm) retinal cross-sections of another cohort of treated or untreated Cre(+) and Cre(–) mice (n = 8 per group) stained with epoxy-tissue stain. Arrows indicate melanosomes in apical processes; horizontal lines highlight areas lacking apical melanosomes. Scale bar, 20 µm. *j, k,* Double-blinded quantification of apical melanosomes across the RPE surface normalized to 100 µm RPE length at 4 weeks (*j*) and 20 weeks (*k*) post-injection. Right panels show paired comparisons of treated versus contralateral untreated eyes. *l, m,* Side-by-side comparison of melanosome localization quantified from experiments using *Myo7b* activation or *MYO7A* supplementation (combined with data from Fig. 4) at 4 weeks (*l*) and 20 weeks (*m*) post-injection. *n,* Quantification of ONL thickness by optical coherence tomography in animals treated with *MYO7A* supplementation (n = 10), *Myo7b* activation (n = 8), and untreated contralateral Cre(+) eyes (n = 18). Statistical analysis: two-tailed paired t test with correction (*j, k,* right panels); one-way ANOVA with Tukey’s post hoc test (*b, c, e, f, l, m, n,* and left panels of *j, k*). Scatter plots show mean ± s.e.m. All source data are provided in the Source Data file.

Four and 20 weeks after *Myo7b* activation, Cre(+) animals exhibited reduced mislocalization of melanosomes in apical regions of the RPE in injected areas compared to uninjected contralateral retinas of the same mice (Fig. 5i). However, the effect was less pronounced in direct comparison with *MYO7A* gene supplementation, as complete restoration of melanosome localization to the level of Cre(-) control animals could not be achieved (Fig. 5j-m). This suggests that MYO7B partially compensates for MYO7A under the conditions used herein.

Compared to *MYO7A* supplementation, *Myo7b* activation led to a slight decrease in ONL thickness of treated retinas (Fig. 5n & Extended Data Fig. 9e-f). Accordingly, while no signs of inflammation or reactive gliosis were detectable in mice and pigs treated with *MYO7A*, a slight increase in GFAP and Iba-1 expression could be observed for mice treated with *Myo7b* (Extended Data Fig. 9g-i).

To analyze global transcriptional changes or potential off-targets after *Myo7b* activation, we performed RNA-seq on the retinas and RPE of subretinally injected Cre(+) mice 20 weeks after injection. Compared to uninjected samples and those from Cre(-) mice, we identified several genes to be differentially regulated (Extended Data Fig. 10a-d). Predicted sgRNA off-target sites overlapped with differentially expressed genes (DEG) transcription start site regions (±2 kb) only when containing at least three mismatches and a one-nucleotide bulge. Such sites are unlikely to support stable sgRNA binding. Thus, the observed DEG signature is more likely to reflect indirect effects of *Myo7b* activation, MYO7B expression, AAV delivery, or the injection procedure than direct off-target activity. Finally, for potential human application, we also identified human-specific sgRNAs that enabled activation of the *MYO7B* gene in HEK293 cells and in human retinal organoids (Extended Data Fig. 11a-e).

Overall, these data suggest that the activated MYO7B protein in the retina and RPE is similarly localized as MYO7A and can at least partially compensate for its function.

## Discussion

In this study, we deliver proof of concept for two different approaches to treating Usher syndrome caused by mutations in the *MYO7A* gene: *MYO7A* supplementation and transcriptional activation of the endogenous *Myo7b* locus using CRISPRa. Due to the relatively high prevalence of USH and the high level of suffering of those affected, these results are of great therapeutic relevance. Recently, the first clinical study in USH1B patients was initiated using *MYO7A* supplementation with dual DNA trans-splicing AAVs^40^ (<u>NCT06591793</u>), but the long-term efficacy results are still pending^61^. By comparison, we have used dual mRNA trans-splicing AAV vectors to supplement the *MYO7A* gene and to deliver the dCas9-VPR CRISPRa module to the murine retina. In the context of *MYO7A* supplementation, we provide initial evidence that mRNA trans-splicing vectors show higher reconstitution efficiency compared to one version of their DNA trans-splicing counterparts which is not identical to those used in the above-mentioned clinical trial.

Our gene supplementation approach did not result in toxic effects in treated eyes in OCT and histological experiments during the period covered by this study. The dual mRNA trans-splicing AAV vector approach is currently being used in the first clinical study for the treatment of Stargardt disease, which is caused by mutations in the *ABCA4* gene, and first signs of visual improvement were reported in treated patients (<u>NCT07002398</u>). Here we provide further evidence that this method is very well suited for the administration of large genes or CRISPR/Cas modules in a mouse model.

The retinal phenotype of the Cre(+) conditional mouse model characterized here is very similar to that described in the naturally occurring Shaker mouse. Cre(+) mice might be more suitable for evaluating retina-specific gene therapies due to the absence of hearing and balance disorders and the associated side effects, or for animal welfare reasons. Notably, we detected residual expression of the MYO7A protein in the retina and in RPE cells in these mice. While the expression of MYO7A in RPE cells is most likely due to the well-documented mosaicism in these cells in the presence of Cre, the residual expression of this protein in the retina is surprising, since a more or less complete deletion of the floxed *Myo7a* region has been detected in this tissue at the genomic and transcript levels. One possible explanation could be that the polyclonal anti MYO7A antibody (Ep1) recognizes a different protein that may be related to MYO7A. Alternatively, there may have been some contamination from RPE cells in the retinal samples. Since MYO7A is expressed more strongly in the mouse RPE compared to photoreceptors, even minor contamination could thus explain a substantial effect in western blot and proteomic experiments.

Although the activation of *Myo7b* clearly improved the localization of melanosomes compared to untreated control eyes, it was less effective than gene supplementation. This could be explained by the assumption that MYO7B is not capable of completely compensating for the function of MYO7A. However, it is also conceivable that the expression levels of *Myo7b* must first be titrated in a dose-finding study to achieve optimal effects. We observed slight toxicity in *Myo7b*-activated retinas, which could reduce the therapeutic benefit. Future dose-finding studies will reveal whether full compensation of the MYO7A function can be achieved by activating *Myo7b* at lower efficiency using lower AAV doses. The activation of homologous genes has already been successfully used in the retina and other tissues in animal models for various genetic diseases^25–27,30,62^. Here, we provide further evidence that this method is, in principle, suitable for the treatment of genetic diseases, underscoring its potential for use in clinical trials.

## Methods

### Cloning

The coding sequence of human *MYO7A* isoform 1 (NM_000260.3) was used to design REVeRT vectors. The sequence was screened for the minimal AG|G split site consensus sequence. Three different split sites were initially tested, and the most efficient was used for all subsequent experiments. Sequences for REVeRT *MYO7A* construct were obtained by PCR amplification of isolated cDNA from HEK293T cells overexpressing *MYO7A* and cloned into previously described REVeRT backbones (pAAV2.1 backbone) using standard cloning techniques^24,31^. The constructs for *Myo7b/MYO7B* transactivation were obtained as previously described^31^. The CMV/GRK1 promoter was replaced with the same version of the short, ubiquitous chicken beta-actin promoter (CAG)^63^ that was also used for REVeRT MYO7A. All constructs were verified by whole plasmid sequencing before use (Eurofins Genomics).

### Cell culture and transfection

HEK293T cells (Takara, 632180) were cultured in Dulbecco’s modified Eagle’s medium (DMEM) GlutaMAX^TM^ (Thermo Fisher Scientific, high glucose formulation) supplemented with 10% FBS (Biochrom) and with 1% penicillin/streptomycin (P/S, Biochrom) at 37 °C in a humidified incubator with 10% CO_2_. The murine retinoblastoma-derived 661W cell line (kindly provided by M. Al-Ubaidi^64^) was maintained in DMEM GlutaMAX^TM^ medium (with pyruvate) supplemented with 10% FBS and 1% Antibiotic-Antimycotic solution (Anti-Anti; Thermo Fisher Scientific) at 37 °C, 5% CO_2_. Immortalized MEF cells were generated as previously described^65,66^,and maintained in DMEM GlutaMAX^TM^ medium supplemented with 10% FBS and 1% P/S at 37 °C, 5% CO_2_. HEK293T cells were transfected using the TurboFect^TM^Transfection Reagent (Thermo Fisher Scientific) according to the manufacturer’s instructions. 661W and MEF cells were transfected with the Xfect^TM^ transfection reagent (Takara) following the manufacturer’s protocol. For co-transfections, plasmids were mixed at equimolar ratios. Cells were harvested 48 h post-transfection for downstream analyses.

### Mice

All animal procedures were performed in accordance with the German Animal Welfare Act (Tierschutzgesetz) and approved by the local regulatory authority (District Government of Upper Bavaria, Germany). Experiments were conducted using C57BL/6J wild-type mice and retina-and RPE-specific *Myo7a* conditional knockout mice (Cre(+)). All mice were housed under standard conditions in a 12-hour light/dark cycle at 22 °C, and 60% humidity with food (Ssniff; maintenance diet: /M-H; breeding diet: M-Z Extrudat) and water provided ad libitum. Euthanasia was performed by cervical dislocation following isoflurane anaesthesia.

### Pigs

All animal procedures were conducted in compliance with Directive 2010/63/EU on the protection of animals used for scientific purposes. The Institute of Animal Physiology and Genetics is authorized to conduct research involving animals under the decision issued by the Ministry of Agriculture of the Czech Republic (File No. MZE-24154/2021-18134, dated April 26, 2021. Surgeries and takedowns of the animals were performed at the PIGMOD Center, Liběchov, Czech Republic.

### Ocular surgery of pigs

Wild-type domestic pigs (*Sus Scrofa domesticus)*, Liběchov minipig strain^67^ aged 5-6 months were used. Both animals received bilateral subretinal injection of dual REVeRT *MYO7A* AAVs using the AAV8(Y733F)^42^ capsid (1×10^12^ total vg in 200 µL) or the AAV2.GL capsid^43^. General anaesthesia was induced via intravenous bolus administration (vena cava cranialis) of propofol 1% (MCT/LCT, Fresenius; 20-30 mL) with rocuronium bromide (10 mg/mL, Hameln; 20-25 mg), followed by tracheal intubation and maintained by continuous intravenous infusion (37 mL/h) of propofol 1% (22 mL) with rocuronium bromide (20 mg). Before surgery, the periorbital region was disinfected with povidone-iodine, and sterile drapes were applied. A lid speculum was inserted, and the 23-G trocars were placed through the pars plana. Core vitrectomy was performed, followed by induction of a localized retinal detachment by subretinal injection of 200 µL vector solution near the visual streak using a 41-G cannula. All surgeries were performed by a single experienced surgeon (MDF). Upon completion, trocars were removed, and dexamethasone (1 mg in 0.5 mL) was administered subconjunctivally. Olfloxacin ointment (0.3%) was applied to the ocular surface before recovery from anesthesia.

### Porcine tissue harvest

38 days after vector administration, animals were examined terminally and euthanized under deep anesthesia. Sedation was induced by intramuscular injection of TKS mixture containing tiletamine (2 mg/kg) and zolazepam (2 mg/kg) (Zoletil 100, Vibrac), ketamine (2 mg/kg) (Narketan 10, Chassot) and xylazine (0.4 mg/kg) (Rometar 2%, Spofa). Anesthesia was deepened with an intravenous bolus (vena auricularis lateralis) of propofol 1% (10 mL) and rocuronium bromide (30 mg) during ocular examination. Immediately before exsanguination, an additional bolus of propofol 1% (10 mL) was administered. Eyes were marked at the nasal sclera for orientation, enucleated, rinsed in PBS, and freed of extraocular muscles. The cornea, lens, and vitreous body were removed. Retinal punches (6 mm and 8 mm) were collected from the injected bleb area and uninjected regions. For immunohistochemistry (IHC), tissue punches were fixed in 4% paraformaldehyde (PFA) for 24 h at 4 °C, dehydrated in ascending sucrose concentrations (10%, 20%, 30%) containing 0.01% sodium azide (each for 2 h at room temperature), frozen in dry ice-cooled isopentane for 1 h, and embedded in tissue-freezing medium (Leica, 14020108926). Cryosections (20-30 µm thick) were obtained using a cryostat (Leica, CM1950). For PCR and western blot analysis, the tissue punches were snap-frozen in liquid nitrogen.

### Human retinal organoid culture and AAV transduction

Human retinal organoids shown in Fig. 2e were generated and cultured according to established protocols^68,69^ in the laboratory of Prof. Dr. Jan Wijnholds at Leiden University Medical Centre. Organoids were maintained at 37 °C in a humidified incubator with 5% CO_2_. All other organoids shown were generated and cultured in-house following the protocol by Kim et al. 2019^70^ and as previously described^24^. Transduction of human retinal organoids was performed at differentiation day 220 (DD220) as previously described^24^, corresponding to a mature differentiation stage. Briefly, dual AAV vectors carrying the expression construct were diluted in 50 µL of organoid culture medium to a final concentration of 1×10^11^ to 1×10^12^ viral genomes per organoid. Each organoid was placed in an individual well of an agarose-coated 96-well plate, and 50 µL of the AAV solution was added directly to the organoid. Organoids were incubated with the AAV solution for 8 hours at 37 °C and 5% CO2, after which 150 µL of fresh culture medium was added to each well. Medium was exchanged every 2-3 days, and organoids were harvested three weeks post-transduction for downstream expression analyses. The AAV2.NN^43^ capsid variant was used for all transduction experiments.

### Generation of Rx-Cre(+)-Myo7a mice

The mouse strain used for this research project, B6(129S4)-*Myo7a^tm1c(EUCOMM)Wtsi^*/SboyeMmmh, RRID: MMRRC_067354-MU, was obtained from the Mutant Mouse Resource and Research Center (MMRRC) at the University of Missouri, an NIH-funded strain repository, and was donated to the MMRRC by Shannon Boye, Ph.D., University of Florida, School of Medicine. These mice carry loxP sequences around essential exons of *Myo7a*^47,71,72^. Conditional *Myo7a* KO mice were generated by crossing *Myo7a* flox/flox mice with a Cre-deleter line expressing Cre recombinase under the control of the *Rax* promoter (retina and anterior neural fold homebox). This line, designated *Rx*-Cre (Bl6.Cg-Tg(Rax-cre)1Zcoz/Ph), was obtained under a material transfer agreement from the Institute of Molecular Genetics of the ASCR, Prague, Czech Republic, and maintained within our animal facility. To obtain the experimental cohort, homozygous *Myo7a* flox/flox (Cre(-)) mice were bred with homozygous *Myo7a* flox/flox (Cre(+)) mice. This breeding strategy yields offspring with a 1:1 ratio of Cre(+) (experimental group) and Cre(-) (wild-type control group) homozygous floxed mice. All animals were genotyped before use in experiments. *Myo7a* was genotyped by using the following primer set:

*Myo7a* loxP forward: 5’TACACATGGGCAATCTGCAG3’ and

*Myo7a* loxP reverse: 5’ACACGCATCCAAGTTCTCAA3’. A band at 941 bp was identified as wild-type, while a band at 1121 bp corresponds to the flox cassette. The presence of Cre-recombinase was confirmed by a 320 bp band after amplification using the following primer set: *Cre* forward: 5’AGCACCAAAGCTCCAGTTACC3’ and *Cre* reverse: 5’CGTTGCATCGACCGGTAATGCA3’.

### Electroretinography (ERG)

Mice were dark-adapted overnight before recordings. For pupil dilation, tropicamide eye drops (Mydriaticum Stulln, Pharma Stulln GmbH) were applied shortly before the procedure. Full-field ERG was performed using a Celeris apparatus (Diagnosys LLC) with integrated light guide electrodes positioned on each cornea. Scotopic ERG responses were recorded following single-flash stimuli of increasing light intensities (0.003, 0.01, 0.03, 0.1, 0.3, 1, 3, 10 cd.s/m^2^). After a 5-minute light adaptation period at 9 cd/m^2^, photopic single-flash responses were recorded using the same series of intensities under continuous 9 cd/m^2^ background illumination.

### Optical coherence tomography (OCT)

Retinal imaging was performed using a modified Spectralis HRA + OCT system (Heidelberg Engineering) equipped with optic lenses and operated via the Heidelberg Eye Explorer software (version 1.10.4.0). Mice were anesthetized and the pupil dilated as described above. 31 OCT scans were acquired in linear scan mode. The outer nuclear layer (ONL) thickness was defined as the distance encompassing the nuclei of rod and cone photoreceptors. For characterization studies, the ONL thickness was calculated from single values measured from the scan going through the optic nerve. For subretinal-injected eyes, the mean ONL thickness was calculated from single values measured across 15 different scans corresponding to the injected area. The same area of the contralateral uninjected eye was analyzed.

### AAV production and subretinal/intravitreal injections of mice

REVeRT dual AAV vectors were produced using either the AAV8(Y733F)^42^, the AAV2.GL or AAV2.NN capsid^43^ as described previously^24^. For subretinal and intravitreal injections, wild-type (WT), *and* Cre(+) mice were anaesthetized by intraperitoneal injection of 0.02 mg/g body weight ketamine and 0.04 mg/g body weight xylazine, and the pupils were dilated using 1% atropine and 0.5% tropicamide eyedrops. Subretinal and intravitreal injections were performed with a 5 µL syringe (Hamilton) and a 34 G needle with a blunt end (Hamilton) and a micromanipulator as previously described^24,73^. WT mice were subretinally injected at P21 with 1 µL of dual titer-matched AAV8(Y733F) REVeRT *MYO7A* vectors. Reconstitution was analyzed 4 weeks after subretinal injection. Control eyes were either injected with only one half of the *MYO7A* AAVs (5’ or 3’*MYO7A*) or with the AAV formulation buffer or remained uninjected. For gene supplementation treatment, one eye of Cre(+) animals was subretinally injected at P21 with 1 µL of dual titer-matched AAV8(Y733F) REVeRT *MYO7A* vectors (1.0×10^10^ vg/µL) and the contralateral eye remained uninjected. For treatment via *Myo7b* activation, one eye of Cre(+) animals was subretinally injected at P21 with 1 µL of dual titer-matched AAV8(Y733F) encoding split dCas9-VPR (1.0×10^10^ vg/µL) and the contralateral eye remained uninjected. Eyes were harvested either 4 or 20 weeks post-injection. To compare intravitreal injection of REVeRT *MYO7A* with subretinal injection, one eye of Cre(+) animals was either subretinally or intravitreally injected at P21 with 1 µL of dual titer-matched AAV2.GL REVeRT *MYO7A* vectors (1.0×10^10^ vg/µL) and the contralateral eye remained uninjected. Eyes were harvested 4 weeks post-injection for RNA and protein extraction.

### RNA isolation and cDNA synthesis

Transfected cells were harvested 48 h after transfection by removing the culture medium and adding 350 µL of the RNA lysis buffer from the peqGOLD Total RNA kit (VWR) supplemented with 20 µL/mL β-mercaptoethanol to each well of the 6-well plate used for transfection. The cells were transferred to a shaking platform (VWR) for 5 min to ensure uniform distribution of the lysis buffer. The lysed cells were transferred to Safe-Lock tubes (Eppendorf). A stainless-steel ball (Retsch) was added to each tube, and the cells were homogenized and disrupted using a mixer mill (Retsch) at 30 Hz for 1 min, followed by 5 min of centrifugation at 21,000 × *g*. The RNA was isolated using the peqGOLD Total RNA kit (VWR) according to the manufacturer’s instructions. For RNA isolation of mouse tissue, eyes were enucleated using surgical scissors and transferred to 35 x 10 mm petri dishes (Sarstedt) containing filter paper soaked in ice-cold 0.1 M phosphate buffer (PB). The eye was punctured at the ora serrata using a 21-G cannula (VWR). Under a stereomicroscope (Stemi 2000, Zeiss), the ora serrata was incised with small surgical spring scissors (Fine Science Tools) to remove the lens, cornea, and vitreous body. For whole eyecup RNA isolation, the entire eyecup was placed in a Safe-Lock tube and snap-frozen in liquid nitrogen. For the separation of retina and RPE, a small incision was made in the eyecups, and the retina was gently detached from the RPE using two fine forceps (Fine Science Tools). Retina and RPE from the same eye were collected into separate Safe-Lock tubes and snap-frozen in liquid nitrogen. For each tube, 350 µL of RNA lysis buffer was added, followed by the addition of one stainless-steel ball (Retsch). The tissue was disrupted twice for 1 min at 30 Hz using the mixer mill MM400 (Retsch). The stainless-steel balls were removed, and the lysates were centrifuged at 21,000 x g for 5 min. RNA was isolated using the peqGOLD Total RNA Kit (VWR) according to the manufacturer’s protocol. For RNA isolation from transduced human retinal organoids, two organoids were pooled in a Safe-lock tube and lysed in 400 µL TRI Reagent (Zymo). Organoids were disrupted with a stainless-steel ball in a mixer mill for 1 min at 30 Hz. The RNA was purified using the Direct-zol^TM^ RNA Microprep Kit (Zymo) following the manufacturer’s instructions. For pig samples shown in Fig. 2k-n RNA and protein was simultaneously isolated using the NucleoSpin TriPrep Kit (VWR) according to the manufacturer’s instructions. Total RNA concentrations were measured using the Nanodrop^TM^ 2000c spectrophotometer (Thermo Fisher Scientific). Complementary DNA (cDNA) was synthesized using the RevertAid First Strand cDNA Synthesis Kit (Thermo Fisher Scientific) for up to 1 μg total RNA.

### Simultaneous gDNA and RNA isolation

For experiments shown in Extended Data Fig. 3c,g,m, gDNA and RNA were isolated simultaneously for each sample. The tissue was isolated as described in the previous section and lysed using 350 µL of RLT buffer (Qiagen) supplemented with 10 µL/mL β-mercaptoethanol. After adding a stainless-steel ball and disrupting the mixture using a mixer mill at 30 Hz for 1 min, the gDNA was isolated using the Zymo-Spin IIC-XL columns (Zymo). Lysates were centrifuged at 5,000 x g for 10 min at 4 °C, and the supernatant was transferred to the extraction columns, followed by centrifugation at 1,500 x *g* for 4 min and subsequently at 10,000 x *g* for 1 min at room temperature. The flow-through was used for RNA isolation using the peqGOLD Total RNA kit (VWR) according to the manufacturer’s instructions. 400 µL of Genomic lysis buffer (Zymo) supplemented with RNase A (Sigma-Aldrich; 1:5,000) was added to the column. This was incubated for 10 min at room temperature, followed by centrifugation for 2 min at 10,000 x *g*. 400 µL of DNA pre-wash buffer (Zymo) and 600 µL of gDNA wash buffer were applied sequentially, with intermediate centrifugation for 1 min at 10,000 x g. A final wash step was performed with 600 µL of gDNA wash buffer, followed by centrifugation for 1 min at 10,000 x *g*. For elution, 100 µL nuclease-free water was added to the column, incubated for 20 min at room temperature, and centrifuged for 2 min at 10,000 x *g*. gDNA concentration was quantified using a NanoDropTM 2000c spectrophotometer (Thermo Fisher Scientific). Isolated gDNA was used for PCR amplification using the Q5 High-Fidelity polymerase (New England Biolabs). All used primers are listed in Supplementary Table 1.

#### RT-qPCR

For reverse transcription PCR (RT-PCR), Q5 High-Fidelity polymerase (New England Biolabs) was used. Real-time quantitative RT-PCR (RT-qPCR) was performed with duplicates on a MicroAmp™ Fast Optical 96-Well Reaction Plate (Thermo Fisher Scientific) using the QuantStudioTM 5 Real-Time PCR system (Thermo Fisher Scientific) and the PowerUpTM SYBR Green Master Mix (Thermo Fisher Scientific). The expression level of all genes was normalized to either murine *Alas*, human *ALAS* or porcine *B2M* and calculated with the 2^-ΔΔC(T)^ method. All primers are listed in Supplementary Table 1. The data was analyzed using the QuantStudioTM Design & Analysis software (Thermo Fisher Scientific).

### Protein extraction

Western blots of murine tissue were either performed using the entire eyecup or the retina and RPE (including choroid) alone. All samples were lysed in 50 µL of 1x RIPA lysis buffer (Merck), which was supplemented with 10% glycerol and a cOmplete^TM^ ULTRA Protease Inhibitor Cocktail tablet (Roche) in 9 mL of H_2_O. All samples were disrupted using a stainless-steel ball in a mixer mill MM400 (Retsch) at 30 Hz for 30 seconds. The lysates were rotated end-over-end (VWR tube rotator) for 60 min at 4 °C and then centrifuged. The protein-containing supernatant was transferred into new tubes, and the protein concentration was determined using the Pierce^TM^ Bradford Protein Assay Kit (Thermo Fisher Scientific) according to the manufacturer’s instructions. The samples were stored at-80 °C until further use. For protein extraction from porcine retinal punches in Fig. 2g-i, 200 µL of 1x RIPA lysis buffer was added to each sample. Tissue disruption was performed in the same way as for murine tissue. After rotation on a tube rotator (VWR), the samples were additionally sonicated three times using a Branson SONIFIER W-450 D (MARSHALL Scientific) at 30% amplitude for a total of 10 pulses (0.3 s pulse on, 0.7 s pulse off). The samples were kept on ice during the sonication process. Further steps, including the protein quantification, were performed in the same way as for the murine tissue. To isolate proteins from human retinal organoids, the organoids were removed from the culture medium, and five organoids were pooled per condition in a Safe-Lock tube. 100 µL of RIPA lysis buffer was added to the pooled organoids, and the same protocol as described for murine tissue was conducted. For the protease cleavage shown in Extended Data Fig. 3, the protein lysate was prepared using RIPA lysis buffer without protease inhibitors, followed by 72 h incubation at 37 °C to activate endogenous proteases.

### Western blotting

For western blotting, 30 – 80 µg of the lysates were incubated in 1x Laemmli sample buffer containing DTT at 99 °C for 5 min. The proteins were separated on an SDS-polyacrylamide gel via gel electrophoresis. GAPDH (1:2000, 14C10, Cell Signaling) and β-actin (1:3000, AC-15, Sigma-Aldrich) were stained for 1 h at room temperature. Myc (1:500, 4A6, Sigma-Aldrich) and the human MYO7A (1:375, EPR7498, Abcam) antibodies that detect the *C*-terminus of MYO7A were incubated overnight at 4 °C. The polyclonal rabbit-Myo7a antibody (1:1000, PTS-25-6790, Proteus Biosciences) (Epitope1) that detects the *N*-terminus of REVeRT MYO7A was incubated for 1 h at room temperature to stain endogenous mouse and human MYO7A. The rabbit-MYO7A (Epitope2) antibody was kindly provided and generated by Prof. Hermann Ammer, LMU Munich, and was incubated overnight at 4 °C (1:500). Human and murine MYO7B were stained overnight at 4 °C with custom-made rabbit antibodies specific to each (1:500 dilution) (kindly provided and generated by Prof. Hermann Ammer, LMU Munich). Secondary antibodies (mouse anti-rabbit IgG-HRP, 1:2000, Santa Cruz Biotechnology) were incubated for 1 h at RT. All antibodies used for western blotting have been validated by the manufacturer or by testing on positive controls. The western blots were imaged, and the relative band intensities were quantified using ImageLab software (Bio-Rad, v5).

### Mass spectrometry-based proteomics

All samples were prepared in 96-well plate format using the optimized SP3 protocol and UHPLC MS grade solvents^74^. The cell pellets were lysed in 1% NP40, 0.2% (w/v) SDS in 25 mM HEPES, pH 7.5. Protein concentration in lysates was determined using bicinchoninic acid assay (BCA) from Pierce (23221). The protein amount was adjusted to 10 μg in a total volume of 10 μL of lysis buffer. The protein was loaded onto a mixture of 1:1 hydrophilic (Sera-Mag SpeedBeads Carboxylate, GE Healthcare, 45152105050350) and hydrophobic carboxylate-coated magnetic beads (Sera-Mag SpeedBeads Carboxylate, GE Healthcare, 65152105050350, 1 μL each at 10 μg/μL) that had been pre-washed three times with 100 μL of H_2_O. The magnetic beads with the protein sample were mixed at 850 rpm for 1 min at room temperature (RT). To initiate the binding, 20 µL of acetonitrile containing 0.25% formic acid was added, and the mixture was incubated at RT for 8 min at 850 rpm. Subsequently, the beads were washed three times with 180 μL of 80% (v/v) EtOH and once with 180 μL acetonitrile, with incubation at RT for 30 s and 850 rpm between each wash. After the last wash, the beads were resuspended in 21 μL of 100 mM ammonium acetate buffer (ABC). The wash steps and ABC buffer addition were performed by a liquid handling robot (Hamilton Microlab Prep). The on-beads digestion was performed with trypsin (Promega, V5113, 1 μg) overnight at 37 °C and 850 rpm. The resulting peptide mixture was eluted from the magnetic beads into a new 1.5 mL tube. The peptides were further eluted (three times) from magnetic beads by adding 25 μL H_2_O, incubated at 40 °C, 850 rpm for 5 min. These three fractions were added to the first elution fraction and acidified by adding 11.4 μL of formic acid (FA, TCI, F0654). The combined fractions were further purified from the remaining magnetic beads on a magnet and transferred into MS vials.

MS measurements were performed on an Orbitrap Eclipse Tribrid Mass Spectrometer (Thermo Fisher Scientific) coupled to an UltiMate 3000 Nano-HPLC (Thermo Fisher Scientific) via a nanospray Flex ion source (Thermo Fisher Scientific) equipped with a column oven (Sonation) and a FAIMS interface (Thermo Fisher Scientific). Peptides were loaded on an Acclaim PepMap 100 μ-precolumn cartridge (5 μm, 100 Å, 300 μm ID x 5 mm, Thermo Fisher Scientific) and separated at 40°C on a PicoTip emitter (noncoated, 15 cm, 75 μm ID, 8 μm tip, New Objective) that was *in-house* packed with Reprosil-Pur 120 C18-AQ material (1.9 μm, 150 Å, Dr. A. Maisch GmbH). Buffer composition. Buffer A consists of MS-grade H_2_O supplemented with 0.1% FA. Buffer B consists of acetonitrile supplemented with 0.1% FA. The LC gradient, from 4 to 35.2% buffer B in 36 min, was used. The flow rate was 0.3 µL/min.

### Data-independent acquisition

The DIA duty cycle consisted of one MS1 scan followed by 30 MS2 scans with an isolation window of the 4 m/z range, overlapping with an adjacent window at the 2 m/z range. An MS1 scan was conducted using the Orbitrap at a resolution power of 60,000 and a scan range of 200–1800 m/z, with an adjusted RF lens at 30%. MS2 scans were conducted using the Orbitrap at a resolution power of 30000, and the RF lens was set to 30%. The precursor mass window was restricted to a 500 – 740 m/z range. HCD fragmentation was enabled as an activation method with a fixed collision energy of 35%. FAIMS was performed with one CV at-45V for both MS1 and MS2 scans during the duty cycle.

Standalone DIA-NN software under version 2.0 was used for protein identification and quantification^75^. First, a spectral library was predicted *in silico* by the software’s deep learning-based spectra, RTs and IMs prediction using Uniprot *Mus musculus* FASTA (containing canonical and isoforms). DIA-NN search settings: FASTA digest for library-free search/library generation option was enabled, together with a match between runs (MBR) option and precursor FDR level set at 1%. The mass accuracy and the scan window were set to 0 to allow the software to identify optimal conditions. The precursor m/z range was changed to 500-740 m/z to fit the measuring parameters.

### Proteome analysis

Perseus (1.6.10.43) was used to log_2_ transform LFQ intensities^76^. Samples were grouped by genotype and filtered according to the detected proteins with 70% detection in at least one group. Missing values were replaced from a normal distribution according to the default settings (width = 0.3, down shift = 1.8), and volcano plots are constructed performing pairwise group comparisons (FDR = 0.05, two-sided t-test, randomizations = 250) in Perseus. Resulting comparisons are then analyzed for significance using the threshold of p. adjusted value < 0.05 (visual cut-off: log10(0.05) = 1.3) and absolute difference > 0.58 (fold change > 1.5)^76^. STRING database (https://string-db.org) is used for pathway enrichment analysis using significantly differentially expressed proteins shown in the volcano plots^77^.

### Immunohistochemistry (IHC) and Confocal Imaging

Mouse eyes were removed and placed in 0.1 M PB. The eyeball was punctured at the ora serrata with a 21-G cannula and fixed in 4% PFA (w/v) for 5 min. The cornea, lens, and vitreous body were removed using a stereomicroscope. The eye cup was fixed in 4% PFA for 15 min at RT, cryopreserved in a 30% sucrose solution (w/v), and cryosectioned into 14 µm slices after embedding in Tissue-Tek O.C.T. Compound (Sakura). For IHC, the retina was protected from light during the whole procedure. For IHC of human retinal organoids, the organoids were removed from the culture medium and fixed in 4% PFA for 15 min, cryopreserved in a 30% sucrose solution, and cryosectioned into 14 µm slices after embedding in Tissue-Tek O.C.T. Compound (Sakura). For different injections, the retinal sections were stained with different antibodies: rabbit anti-PDE6B (1:2000, PA1-722, Thermo Fisher Scientific) and mouse anti–rhodopsin 1D4 (1:1500, 1D4, Merck) served as markers for rod outer segments, and rabbit anti-ARL13B (1:500, 17711-1-AP, Proteintech) was used as a primary cilia marker. The mouse anti-Myc-tag antibody (1:200, 9B11, Cell Signaling) and the rabbit anti-Myo7a antibody (1:200, PTS-25-6790, Proteus Biosciences) were used to stain transgenic and endogenous MYO7A. The rabbit anti-MYO7B antibody (1:200, 14467-1-AP, Proteintech) was used to stain MYO7B in transactivated retinal tissue. Guinea pig anti-IBA1 (1:200, 234308, Sigma-Aldrich) and mouse anti-GFAP-Cy3-conjugate (1:1000, C9205, Sigma-Aldrich) were used to check for inflammation and reactive gliosis in injected retinas. After overnight incubation at 4 °C, the following secondary antibodies were incubated for 1.5 h at room temperature: Alexa Fluor 555 anti-rabbit (1:500, Cell Signaling), Alexa Fluor^TM^ 488 anti-mouse IgG2a (1:500, Thermo Fisher Scientific), and Alexa Fluor^TM^ 488 anti-guinea pig (1:400, Thermo Fisher Scientific). All antibodies used for IHC have been validated by the manufacturer. For nuclear staining, DAPI solution (1:1000 in 0.1M PB, Thermo Fisher Scientific) was incubated for 10 min. Images of immunolabeled retinal cryosections were captured using a Leica TCS SP8 spectral confocal laser scanning microscope (Leica Microsystems). Image acquisition was performed at z-stacks using the LAS X software (Leica Microsystems). Multiple z-stack images were merged using maximum projection and further processed with the open-source Fiji/ImageJ software (https://fiji.sc, 1.54f, National Institutes of Health).

### Ultrastructure-Expansion Microscopy (U-ExM)

U-ExM was employed to resolve the subcellular and ciliary localization of proteins in photoreceptor cells. Mouse retinal cryosections (14 µm) were prepared as described above. Tissue processing, including expansion, immunostaining, imaging, and quantitative analysis, was performed following established protocols^24,51^. U-ExM was carried out following the procedure reported by Arsenijevic et al. 2024^78^, with slight adjustments. Briefly, cryosections mounted on slides were thawed at room temperature (RT) for 2 min. A double-sided adhesive spacer (0.3 mm thick; IS317, SunJin Lab Co.) was then placed around the tissue section to form a chamber. Then, sections were incubated with the crosslinking prevention solution composed of 2% acrylamide (AA; Sigma-Aldrich, A4058) and 1.4% formaldehyde (FA; Sigma-Aldrich, F8775) in a final volume of 300 µL at 37 °C for 3 hours or overnight. After removing this solution, the slides were placed on a pre-cooled metal block for the pre-gelation step. A 130 µL pre-gel mixture was added, consisting of 75 µL sodium acrylate (38% w/w stock diluted in nuclease-free water; Sigma-Aldrich, 408220), 37.5 µL AA, 7.5 µL N,N′-methylenbisacrylamide (2%; Sigma-Aldrich, M1533), and 10 µL 10× PBS. After 15 minutes, a second polymerization solution containing the same monomers, supplemented with 0.5% ammonium persulfate (APS; Thermo Fisher, 17874) and 0.5% TEMED (Thermo Fisher, 17919), was applied. A 24-mm coverslip was placed on top to close the spacer chamber. Polymerization proceeded on ice for 15 minutes and then at RT for 2 hours. Following gelation, the coverslip and spacer were carefully removed, and the slide was immersed in 50 mL of denaturation buffer (200 mM SDS, 200 mM NaCl, 50 mM Tris, pH 9) at 95 °C for 2 hours in a water bath. The gels were then washed and expanded through three successive incubations in deionized water before immunostaining.

### Immunostaining of Expanded Gels and Image Acquisition

Expanded gels were first shrunk in three 5-minute washes of 1× PBS. Gels were then incubated overnight at 4 °C with primary antibodies diluted in PBS containing 2% bovine serum albumin (BSA). Anti-alpha-(ABCD antibodies_AA345) and beta-(ABCD antibodies AA344) tubulin antibodies were used in combination (1:250 each). MYO7A was stained using the rabbit anti-Myo7a antibody (1:250, PTS-25-6790, Proteus Biosciences), myc using the mouse anti-Myc-tag antibody (1:200, 9B11, Cell Signaling). The anti-rhodopsin antibody (1:1000, MA5-11741, Thermo Fisher Scientific) was used to stain rhodopsin. After three 5-minute washes in PBS with 0.1% Tween-20 (PBST), secondary antibodies were applied for 3 hours at 37 °C. Following another round of washes (3 × 5 minutes in PBST), gels were fully re-expanded by three 15-minute incubations in deionized water. Imaging was performed with a Leica Thunder DMi8 inverted microscope using either a 20× (0.40 NA) or a 63× (1.4 NA) oil objective. Images were acquired in Thunder SVCC mode (small volume computational clearing) with maximum resolution, adaptive strategy, and water mounting. To minimize gel movement during acquisition, expanded samples were mounted on Poly-D-lysine-coated 24-mm coverslips (Gibco A3890401; Marienfeld 0117640).

### Ultramicrotome sectioning

All mouse eyecups were isolated at the same time of day to avoid bias regarding light adaptation. The eyecups were isolated as described for IHC, marked on the temporal side, and fixed overnight at 4 °C in 2% glutaraldehyde and 2% PFA in 0.1M phosphate buffer. Following fixation, the nasal portion (injected site) of each eyecup (treated and untreated) was processed using standard protocols involving osmium tetroxide post-fixation, graded acetone dehydration, and embedding in Epon Embed 812 (Electron Microscopy Sciences) resin. Semi-thin (0.5 µm) transverse sections were prepared on an Ultracut E ultramicrotome (Reichert-Jung), mounted on slides, and stored at room temperature until staining and imaging.

### Melanosome staining and analysis

Semi-thin retinal sections (0.5 µm) were stained with Epoxy Tissue Stain (Electron Microscopy Sciences) to visualize melanosomes in the RPE, following the manufacturer’s protocol, and mounted in PermaFluor aqueous medium (Thermo Fisher Scientific). Stained sections were imaged under a Zeiss Apotome microscope (40x objective) by a blinded operator. For each section, 10-15 images were acquired to span the complete retinal cross-section. Apical melanosomes were counted by a blinded operator in Fiji/ImageJ (https://fiji.sc, 1.54f, National Institutes of Health) using the multi-point tool, and values were normalized to a 100 µm RPE length.

### RNA-Seq

Total RNA was sent to a commercial provider (Azenta Life Sciences) to perform library preparation and sequencing. Paired-end sequencing was performed with polyA enriched mRNA on an Illumina HiSeq system with > 40 million reads per sample and 150 bp read length. Antisense-oriented, paired-end reads were aligned to the mouse reference genome (GRCm39, GENCODE release M36, basic annotation, excluding alternative contigs) using STAR aligner v2.7.11a^79^.Gene-level counts were generated with the featureCounts function from the Rsubread package v2.18^80^. Differential expression between experimental groups was assessed with edgeR v4.2.1^81^ and limma v3.60.3^82,83^. Gene set enrichment analysis was performed using fgsea v1.30.0^84^. Predicting sgRNA off-target sites was done with Cas-OFFinder v2.4^85^ Cas-OFFinder-bulge v1.2 beta. Differentially expressed genes were defined as those with an absolute log2 fold change > 0.585 and an adjusted p value <0.05

### Statistical analysis and reproducibility

All values are given as mean ± SEM. The number of replicates (n) is stated in the figure legends and can be inferred from scatter plots for each experiment. Statistical analysis was performed using GraphPad PRISM (GraphPad Software, v10.0.2). Normality was tested using a Shapiro-Wilk test. For parametric unpaired data, unpaired t-test with Welch’s correction or One-way-ANOVA was used. For non-parametric unpaired data, Mann-Whitney or Kruskal-Wallis test was used. Multiple comparisons were corrected using Tukey’s or Dunnett’s T3 test. In experiments with paired samples, a paired t-test with Welch’s correction was employed. All RT-qPCR, PCR and western blots have been repeated a minimum of 2 times. All immunohistochemical stainings shown had been repeated for a minimum of 5 more sections. No statistical method was used to predetermine sample size. No data were excluded from the analyses. All animals and cells were randomly assigned to the experimental or the control group. The investigators were not blinded to allocation during experiments and outcome assessment except for the quantification of apical melanosomes. Melanosomes were quantified in a double-blinded manner. First, the sections were imaged blindly, followed by blinded manual quantification of apical melanosomes across the RPE surface. These experiments were repeated 2 times.

## Data availability

All source data are provided as a Source Data file. All omics datasets generated and analyzed in this study (RNAseq and proteomics data) will be made publicly available in a dedicated, publicly accessible repository (e.g. NIH GEO^86^ for next-generation sequencing data and PRIDE^87^ for proteomics data) upon publication in a peer reviewed journal. All primer sequences used in this study can be found in Table 1. The *Mus musculus* GRCm38.p6 and *human* GRCCh38.p13 reference genome can be accessed via Ensembl (http://nov2020.archive.ensembl.org/Mus_musculus/Info/Index) (https://nov2020.archive.ensembl.org/Homo_sapiens/Info/Index).

## Acknowledgements

The authors thank Berit Noack, Kerstin Skokan, and Daniela Lenz for technical support. We thank M. Al-Ubaidi for the gift of the 661 W cells. This work was supported by the Swiss National Science Foundation (310030_212190 and 320030E_221942 to E.B.) and the German Research Foundation (DFG) (project number 513025799 / FOR5621 to E.B., M.B., S.K., and S.M. and project number 325871075 to S.M.). This work has also been supported by the University Research Priority Program of the University of Zurich (URPP) ITINERARE – Innovative Therapies in Rare Diseases (to E.B.) and SAVE SIGHT NOW EUROPE (to Z.E., and E.B.)

