## Supplementary Material for "Gene Supplementation of *MYO7A* or activation of *Myo7b* for treatment of Usher syndrome 1B"

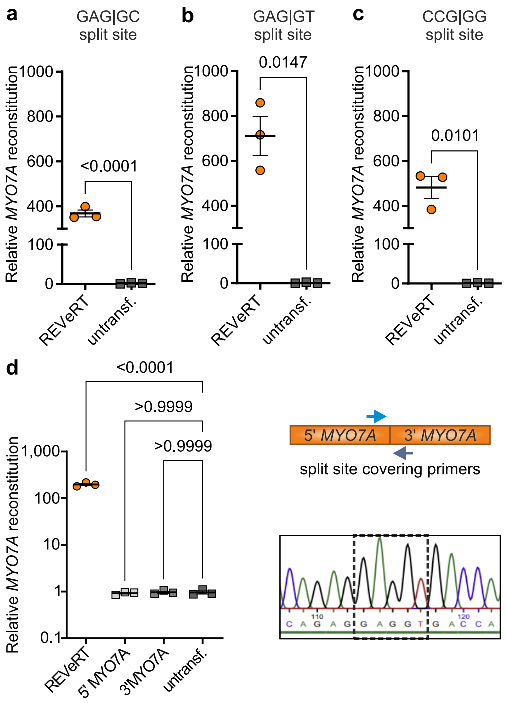


**Extended Data Fig. 1 Validation of *MYO7A* reconstitution *in vitro*.**
*a–c,* RT-qPCR from HEK293T cells transfected with pAAV plasmids co-expressing *MYO7A* constructs split between exons 25/26 (*a*), 26/27 (*b*), and 27/28 (*c*), using split site spanning primers specific for each junction. *d,* Left: RT-qPCR from co-transfected 661W cells with split site spanning primers. Right: Sanger sequencing of reconstituted *MYO7A* from co-transfected MEF cells using split site spanning primers shown in Fig. 2. Statistical analysis, Two-tailed unpaired t test with Welch's correction (*a–c*); one-way ANOVA with Tukey's post hoc test (*d*). Scatter plots show mean ± s.e.m. All source data are provided in the Source Data file.

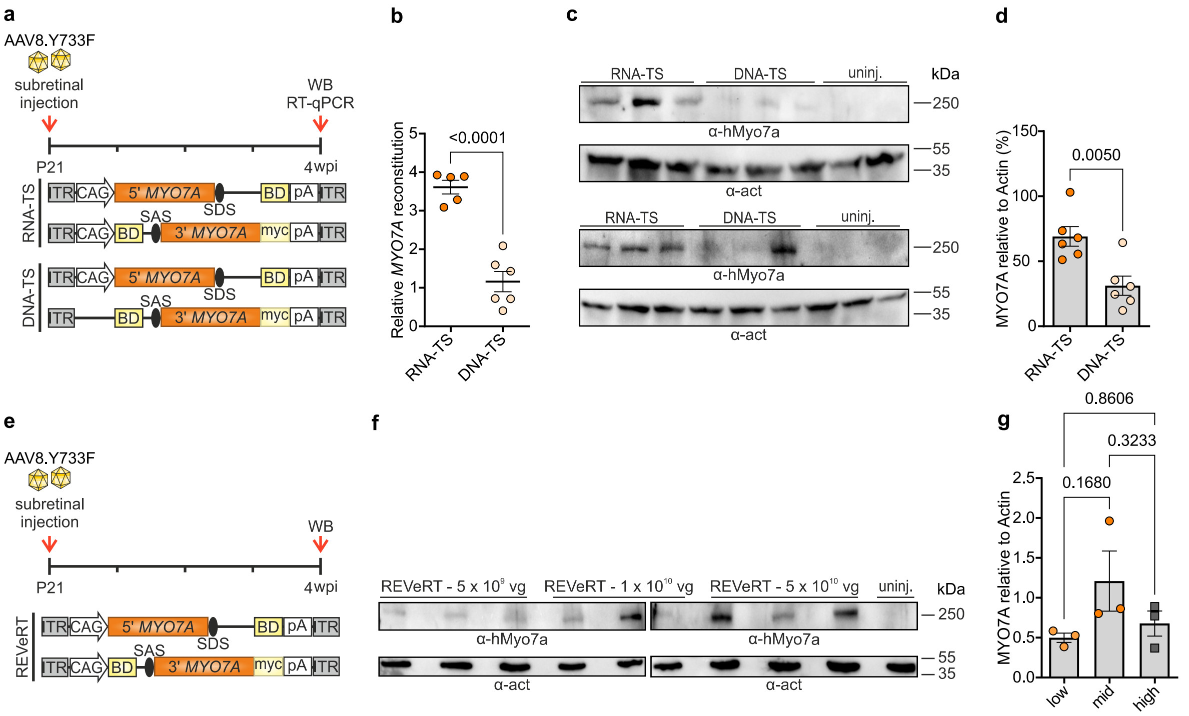


**Extended Data Fig. 2** ***In vivo*** **comparison of different types and doses of dual AAV vectors**

*a,* Overview of the experiments for the side-by-side comparison of mRNA- and DNA-trans-splicing dual AAVs (RNA-TS or DNA-TS). Upper panel, dual AAVs ($1 \times 10^{10}$ total vg; $5 \times 10^{9}$ vg per vector) were subretinally injected to wild-type mice. *b,* RT-qPCR of injected eyecups (RNA-TS, n = 5; DNA-TS, n = 6; uninjected, n = 5) using split site spanning primers to quantify relative *MYO7A* reconstitution (data adapted from Mittas et al. 2026[24]). *c,* Representative western blots of eyecups injected with the respective dual AAVs (n = 6 per injected group, n = 5 uninjected; upper blot n = 1-3, lower blot n = 4-6). *d,* Semi-quantitative analysis of c. *e,* Experimental workflow for dose-testing of dual RNA-TS AAVs expressing *MYO7A*. The vectors were injected to wild-type mice using one of three different AAV doses shown in f. *f,* Representative western blot of eyecups 4 weeks post-injection. *g,* Semi-quantitative analysis of f. Statistical analysis: two-tailed unpaired t test with Welch's correction (*b, d*); and one-way ANOVA with Tukey's post hoc test (g). Scatter plots show mean ± s.e.m. All source data are provided in the Source Data file.

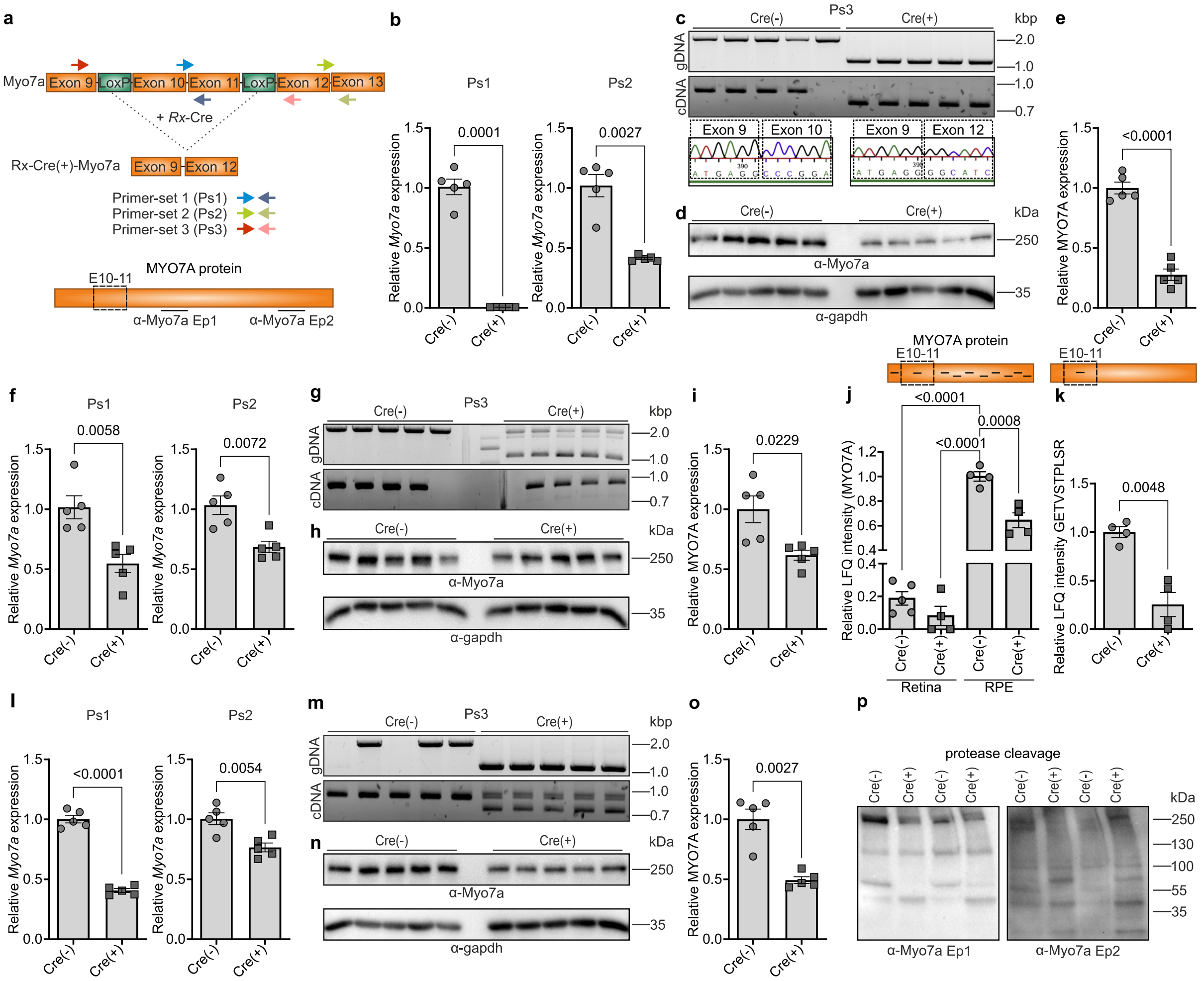


**Extended Data Fig. 3 *Myo7a* expression analysis in the retina and RPE of Cre(+) mice.**
*a,* Schematic of floxed *Myo7a* alleles carrying loxP sites around exons 10 and 11. Homozygous floxed mice (Cre(–)) were crossed with homozygous floxed mice (Cre(+)) to generate Rx-Cre(+) mice (homozygous floxed-Cre(+)) and Cre(–) littermates as controls. Coloured arrowheads indicate the primer-binding sites used *in the* subsequent panels. Epitopes (Ep) for the used MYO7A antibodies (α-Myo7a) are indicated in the scheme of the MYO7A protein. *b,* Relative *Myo7a* expression in neuronal retina of 4 week old Cre(+) and Cre(–) mice (n = 5) using primers inside (PS1) or outside (PS2) the deleted region. *c,* PCR amplification of gDNA and cDNA from neuronal retina with primers spanning the deleted region (PS3) (n = 5). Lower panel shows Sanger sequencing of amplified *Myo7a* transcripts. *d,* Western blot of retinal lysates from Cre(+) and Cre(–) mice (n = 5) probed with a pan-M7A antibody. *e,* Semi-quantitative analysis of MYO7A expression normalized to GAPDH. *f,* Relative *Myo7a* expression in RPE of 4 week old Cre(+) and Cre(–) mice (n = 5) using primers inside (PS1) or outside (PS2) the deleted region. *g,* PCR amplification of gDNA and cDNA from RPE with primers spanning the deleted region (PS3) (n = 5). *h,* Western blot of RPE lysates (n = 5). *i,* Semi-quantitative analysis of MYO7A expression from h, normalized to GAPDH. *j, k,* Proteomics-based quantification of all detected MYO7A peptides covering the entire protein region as indicated by the black bars in the scheme (*j*) and a single peptide corresponding to the deleted region (*k*). *l,* Relative *Myo7a* expression in eyecups of 4 week old Cre(+) and Cre(–) mice (n = 5) using primers inside (PS1) or outside (PS2) the deleted region. *m,* PCR amplification of gDNA and cDNA from eyecups with primers spanning the deleted region (PS3) (n = 5). *n,* Western blot of eyecup lysates from Cre(+) and Cre(–) mice (n = 5) probed with a pan-M7A antibody. *o,* Semi-quantitative analysis of MYO7A levels normalized to GAPDH. *p,* Western blot analysis of RPE lysates (n = 2) after proteasomal cleavage using two different pan-MYO7A antibodies with different epitopes (Ep). Statistical analysis: two-tailed unpaired t test with Welch's correction (*b, e, f, i, j, k, l, o*). Scatter plots show mean ± s.e.m. All source data are provided in the Source Data file.

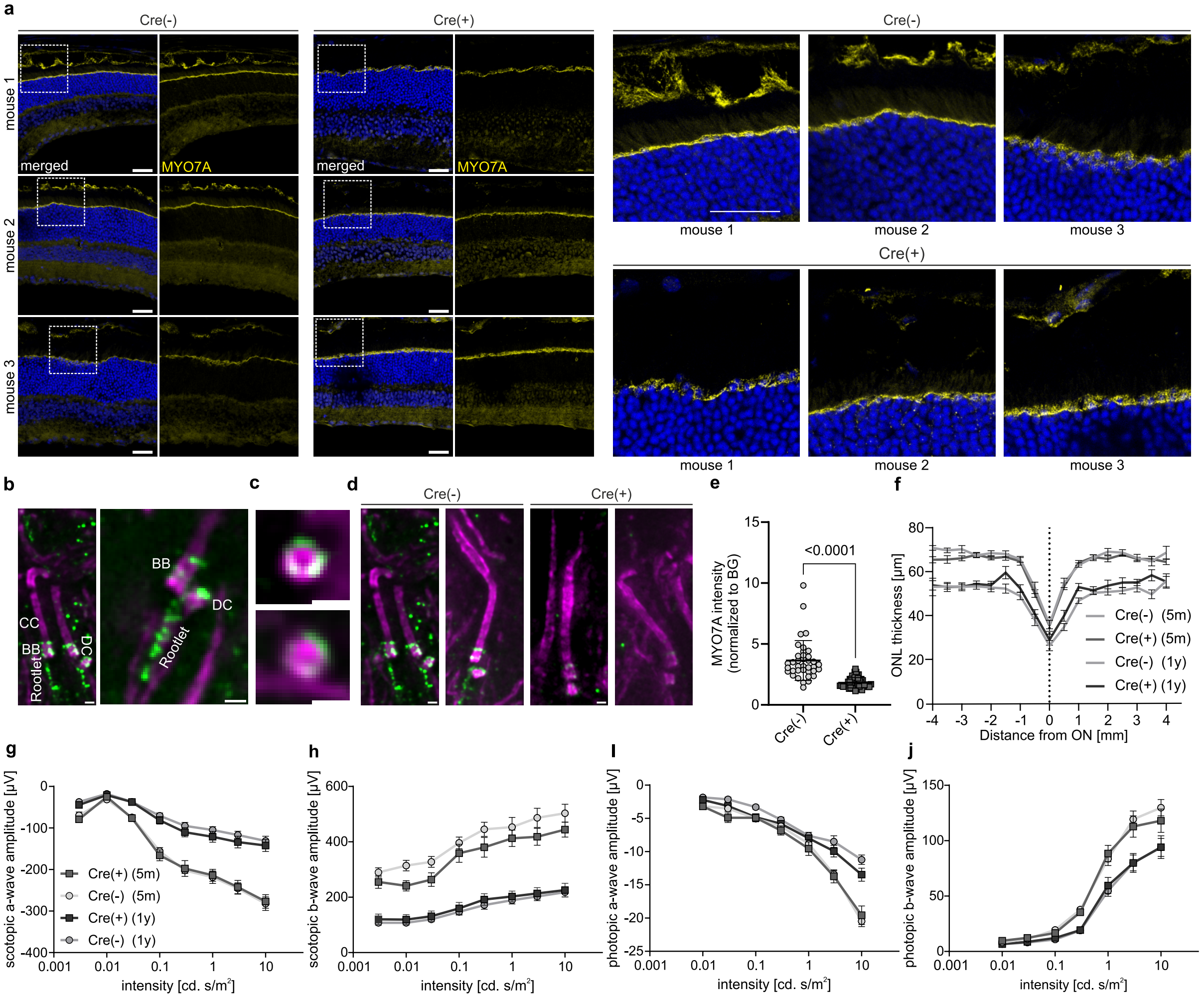


**Extended Data Fig. 4 Retinal morphology and function of Cre(+) mice.**

*a,* Representative immunostaining of retinal cryosections from Cre(+) (n = 3) and Cre(–) (n = 3) mice using a pan-M7A antibody. Arrowheads indicate mosaic MYO7A expression in the RPE of Cre(+) mice. Lower panels show enlarged view. Scale bar, 30 µm. *b,* Representative immunostaining of single photoreceptor cells from Cre(–) mice after expansion. Scale bar, 500 nm. *c,* Transversal section within the transition zone of the ciliary basal body. Scale bar, 50 nm. *d,* Representative immunostaining of single photoreceptor cells from Cre(–) and Cre(+) mice after expansion (n = 2 mice per group). Scale bar, 500 nm. *e,* Quantification of MYO7A intensity at the basal body and transition zone, normalized to background (BG); each datapoint represents a single photoreceptor cell (n = 2 mice/eyes per group). *f,* ONL thickness in relative distance to the optic nerve (ON) measured by OCT in 5 months old (n = 16 per group) and 1 year old (n = 16 per group) Cre(+) and Cre(–) eyes. *g–j,* ERG measurements from 5 months old (n = 10 per group) and 1 year old (n = 10 Cre(+) and 16 Cre(-)) Rx-Cre(+) mice (Cre(+) and Cre(–)) under scotopic (*f–g*) or photopic (*h–i*) conditions. Statistical analysis: two-tailed Mann–Whitney test (*e*). Scatter plots show mean ± s.e.m. All source data are provided in the Source Data file. CC, connecting cilium; BB, basal body; DC, daughter centriole.

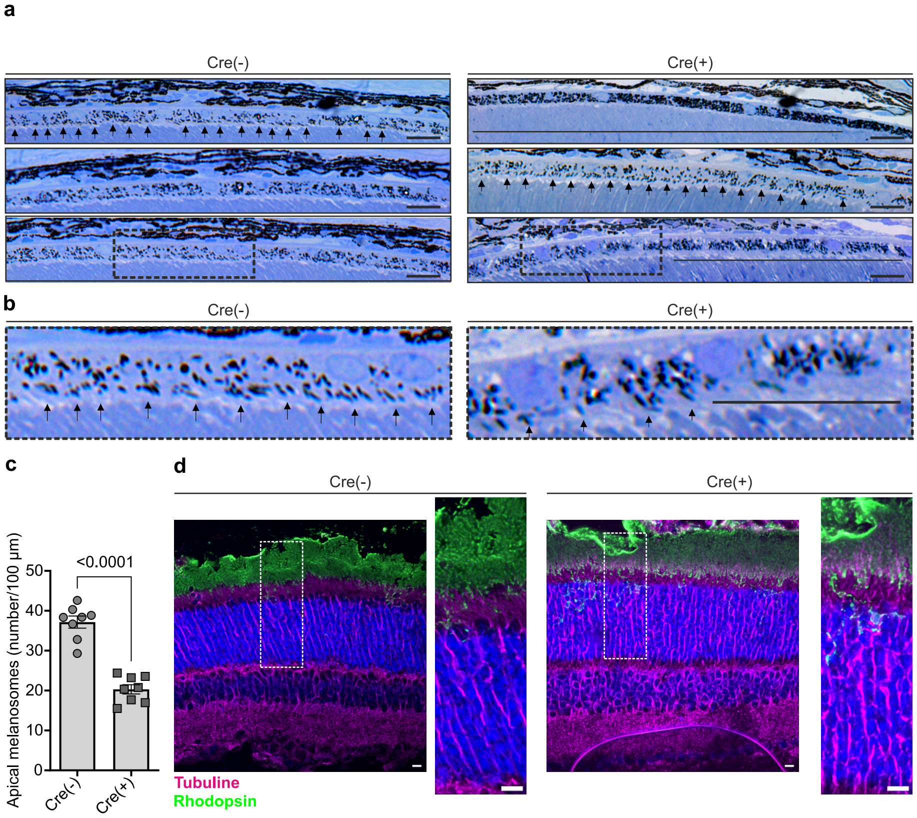


**Extended Data Fig. 5 Phenotypic characterization of Cre(+) mice.**

*a, b,* Representative semi-thin (500 nm) retinal cross-sections of 4 week old Cre(+) and Cre(–) mice (n = 8 per group), stained with epoxy-tissue stain. Arrows indicate exemplary melanosomes in apical processes; horizontal lines highlight areas lacking apical melanosomes. Scale bar, 20 µm. *c,* Quantification of apical melanosomes across the RPE surface, normalized to 100 µm RPE length. *d,* Representative immunostainings of expanded retinas from Cre(+) and Cre(–) mice (n = 2 per group) using ultrastructure expansion microscopy. Right panels show close-up images of the regions marked by dashed rectangles. Scale bar 20 µm., Statistical analysis: two-tailed unpaired t test with Welch's correction (*c*). Scatter plots show mean ± s.e.m. All source data are provided in the Source Data file.

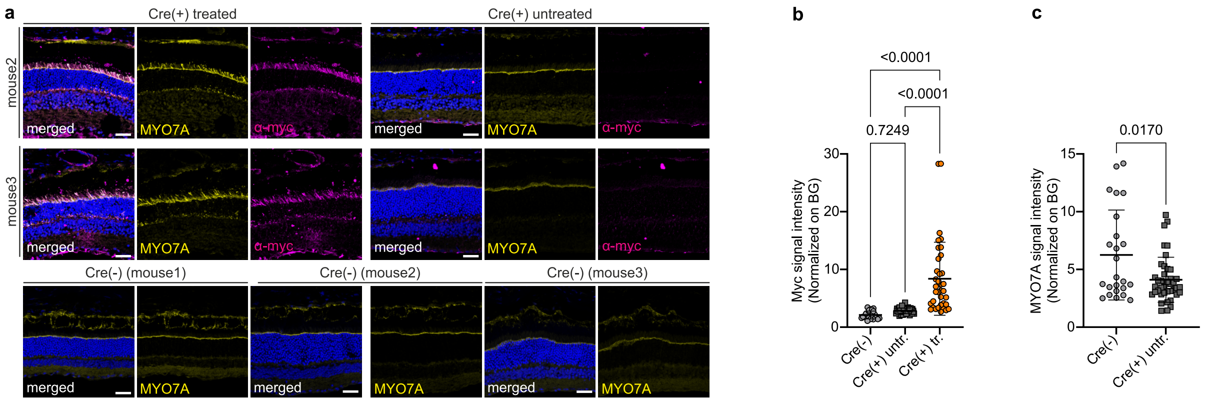


**Extended Data Fig. 6 Localization and expression of MYO7A after gene supplementation to Cre(+) mice.**

*a,* Representative immunostaining of Cre(+) mice 5 months after subretinal injection of dual mRNA trans-splicing AAVs expressing *MYO7A* (includes mouse 2 and 3 from mouse 1 in Fig. 4, and Cre(–) controls). Images show injected and uninjected regions. Scale bar, 30 µm. *b,* Quantification of myc signal intensity at the ciliary basal body, normalized to background, in stained and expanded tissue sections. *c*, Quantification of MYO7A signal intensity (Cre(-) vs. Cre(+) untreated) at the ciliary basal body, normalized to background. Statistical analysis: two-tailed unpaired t test with Welch's correction (*b,c*).

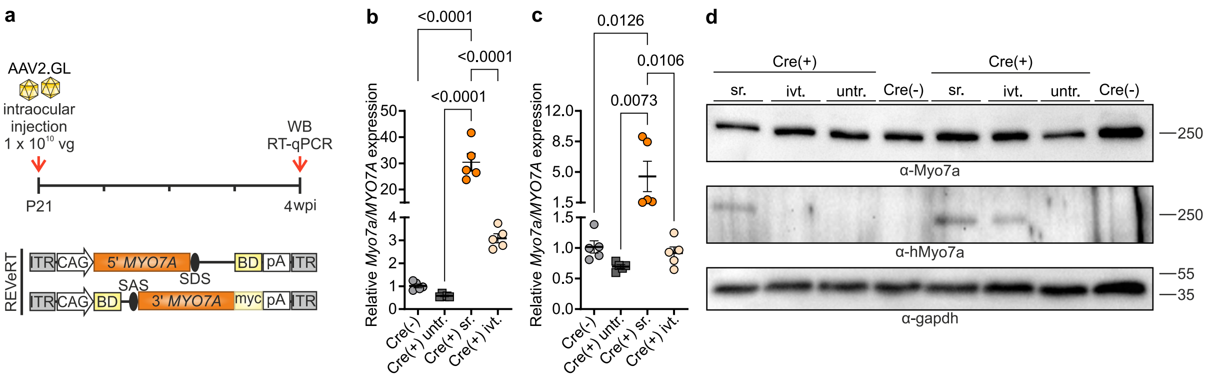


**Extended Data Fig. 7 Reconstitution of *MYO7A* following intravitreal injection**

*a,* Experimental design: one eye of Cre(+) mice was injected subretinally or intravitreally with dual AAVs expressing *MYO7A*, while the contralateral eye remained uninjected. Age-matched Cre(–) mice served as naïve controls. Retinas and RPEs were harvested 4 weeks post-injection. *b, c,* qPCR showing relative *Myo7a*/*MYO7A* expression in neuronal retina (*b*) and RPE (*c*) using split site spanning primers (n = 5 per group). *d,* Representative western blot of retinal lysates from subretinally and intravitreally injected eyes probed with a human-specific or pan-M7A antibody; Cre(+) untreated and Cre(–) eyes served as controls (n = 2 per group). Statistical analysis: one-way ANOVA with Tukey's post hoc test (*b*, *c*). Scatter plots show mean ± s.e.m. All source data are provided in the Source Data file.

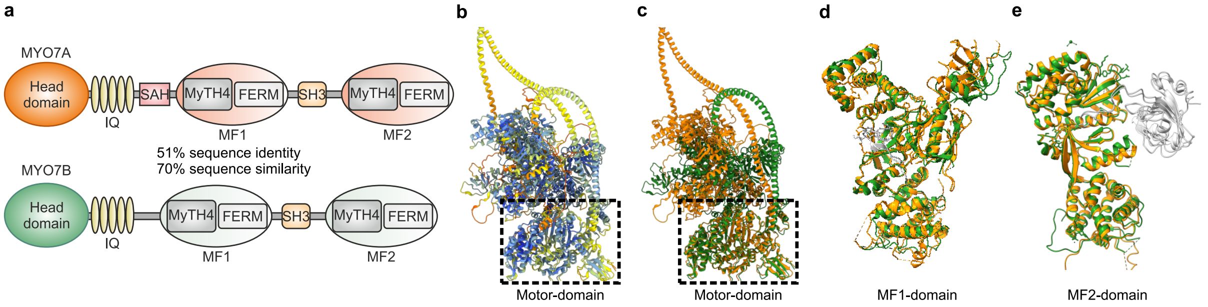


**Extended Data Fig. 8 Structural comparison of MYO7A and MYO7B**

*a*, Domain organization of mouse MYO7A and MYO7B. Both proteins share a conserved architecture consisting of a motor domain, five IQ motifs, and tandem MyTH4-FERM-SH3 modules (MF1 and MF2). Sequence comparison revealed 51.7% identity and 70.4% similarity. *b*, Structural superposition of full-length AlphaFold models colored according to pLDDT confidence scores. High-confidence regions are shown in blue, whereas lower-confidence regions are displayed in cyan, yellow, and orange. *c*, Structural superposition of full-length MYO7A (orange) and MYO7B (green) generated using ChimeraX MatchMaker. Alignment of the conserved core yielded an RMSD of 0.865 Å across 570 pruned atom pairs. *d*, Structural superposition of the N-terminal MyTH4-FERM-SH3 (MF1) modules of human MYO7A and human MYO7B in complex with SANS and ANKS4B, respectively. MYO7A and MYO7B are shown in orange and green, whereas binding partners are displayed in gray. Structural alignment yielded an RMSD of 1.069 Å across 322 pruned atom pairs. *e*, Structural superposition of the C-terminal MyTH4-FERM (MF2) domains of human MYO7A and human MYO7B in complex with harmonin. MYO7A and MYO7B are shown in orange and green, whereas harmonin is displayed in gray. Structural alignment yielded an RMSD of 1.120 Å across 402 pruned atom pairs.

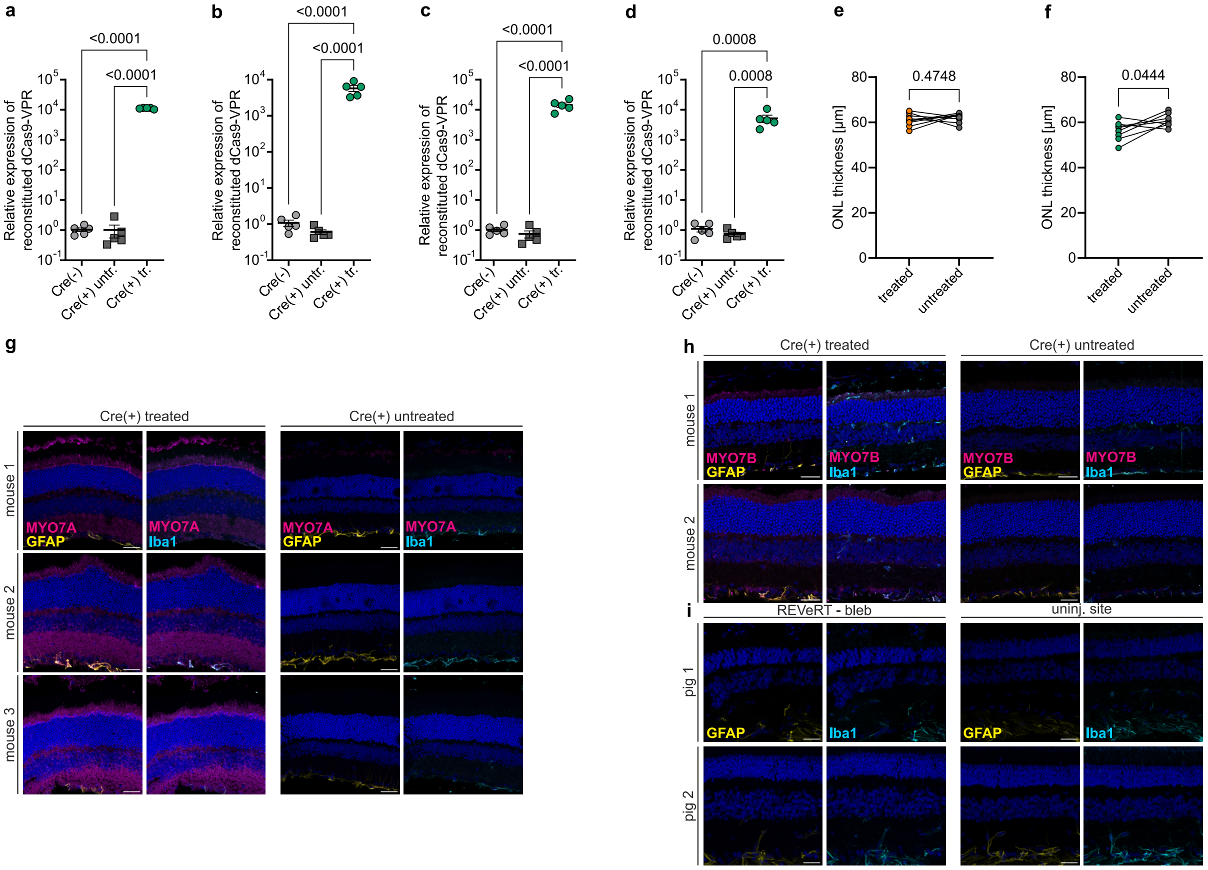


**Extended Data Fig. 9 Assessment of expression, morphology, inflammation and reactive gliosis in injected mice and pigs treated with dual AAVs for *MYO7A* supplementation or *Myo7b* activation**

*a, b,* Relative expression of reconstituted dCas9 in the neuronal retina (*a*) and RPE (*b*) four weeks post-injection (n = 5 per group). *c, d,* Relative expression of reconstituted dCas9 in the neuronal retina (*c*) and RPE (*d*) twenty weeks post-injection (n = 5 per group). *e, f,* Paired comparison of ONL thickness between treated and contralateral untreated eyes per mouse: *MYO7A* gene supplementation-treated eyes (n = 9) in *e* and *Myo7b*-activated eyes (n = 8) in *f. g,* Representative immunostainings of Cre(+) mice injected with dual AAVs expressing *MYO7A*. *h,* Representative immunostainings of Cre(+) mice subretinally injected with titer-matched dual dCas9–VPR-expressing AAVs ($1x10^{10}$ vg in total) targeting the *Myo7b* locus. *i,* Representative immunostainings of wild-type mini pigs subretinally injected with titer-matched dual AAVs ($1x10^{12}$ vg in total) expressing *MYO7A* AAVs. Scale bar, 30 µm.

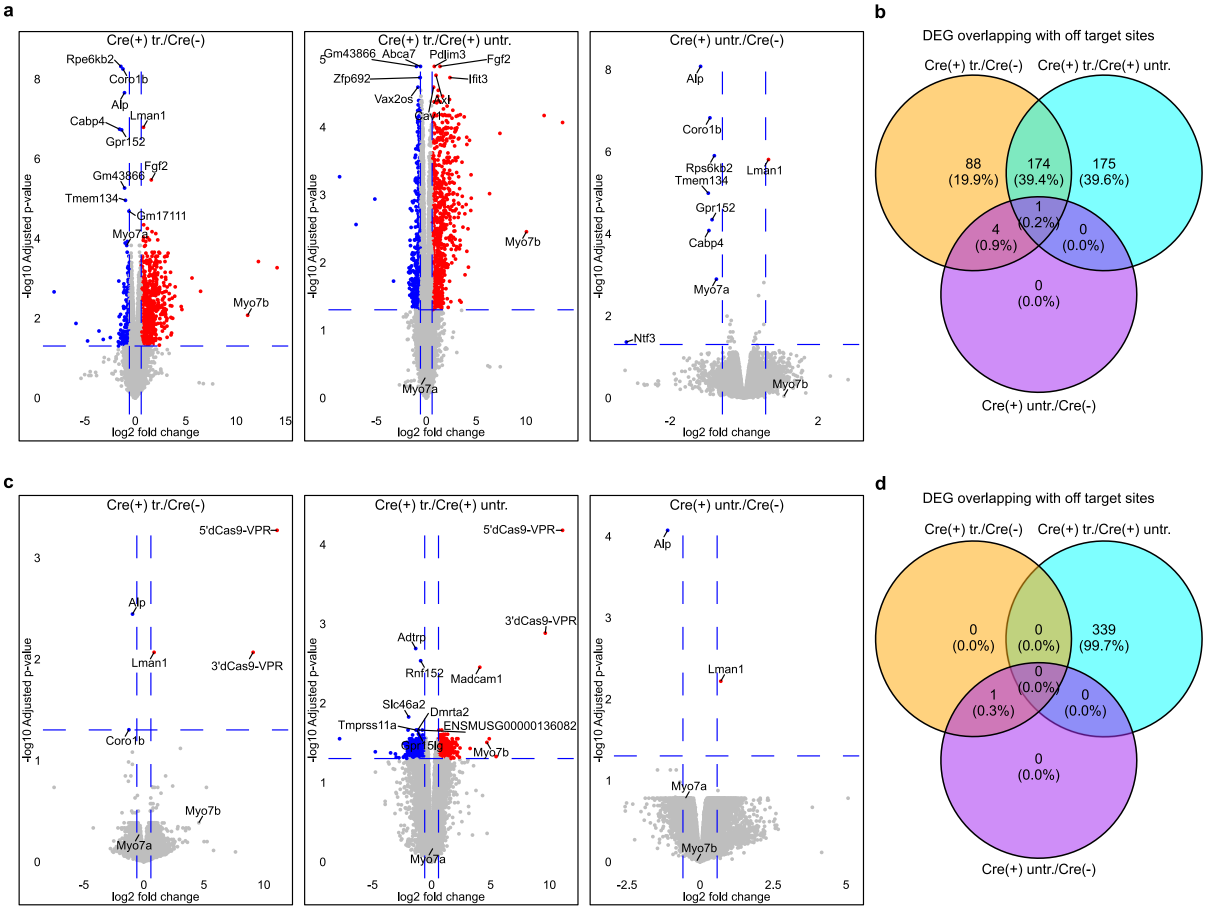


**Extended Data Fig. 10 Off-target analyses in Cre(+) mice treated with dual AAVs for *Myo7b* activation**

*a-d,* Volcano plots showing pairwise comparisons between treatment groups in the retina (*a*) and retinal pigment epithelium (RPE) (*c*). DEGs were defined by |logFC| ≥ 0.585 and an adjusted p-value < 0.05. Downregulated genes are indicated in blue and upregulated genes in red. The indications 5'dCas9-VPR and 3'dCas9-VPR indicate expression of the respective dCas9-VPR transcript half. DEGs overlapping predicted sgRNA off-target sites close to promoter regions (+-2000 nt from TSS) are shown as Venn diagrams for the retina (*b*) and RPE (*d*). Statistical analysis: two-tailed paired t test (*e, f*); one-way ANOVA with Tukey's post hoc test (*a–d*). Scatter plots show mean ± s.e.m. All source data are provided in the Source Data file.

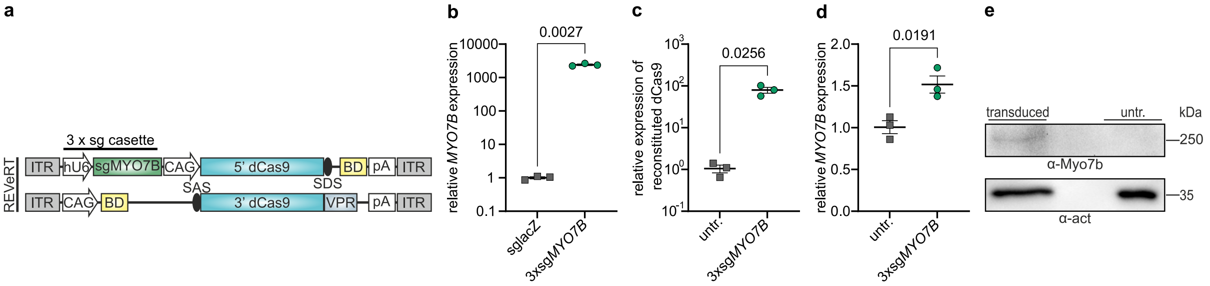


**Extended Data Fig. 11 Human *MYO7B* activation in HEK293 cells and human retinal organoids**
*a,* Schematic of the dual AAV vector expression cassette used to activate the human *MYO7B* locus with sgRNAs targeting its promoter. *b, c,* Relative transcript expression of *MYO7B* (*b*) and reconstituted dCas9 (*c*) following cotransfection of HEK293 cells with the REVeRT plasmids shown in a. *d,* Relative transcript expression of *MYO7B* three weeks after cotransduction of mature wild-type human retinal organoids with titer-matched dual dCas9–VPR expressing AAVs (n = 3 per group). *e,* Representative western blot of transduced retinal organoids (total dose 5 × $10^{10}$ vg; n = 5 organoids pooled per group). Statistical analysis: two-tailed unpaired t test with Welch's correction (*b*–*d*). Scatter plots show mean ± s.e.m. All source data are provided in the Source Data file.

**Supplementary Table 1:** Primer sequences used

| Name | Sequence (5' → 3') |
|---|---|
| qPCR mouse Alas fw | CAGGAGGACGTGCAGGAAAT |
| qPCR mouse Alas rev | CGGCTTGGATCCTCTCCATC |
| qPCR human ALAS fw | GATGTCAGCCACCTCAGAGAAC |
| qPCR human ALAS rev | CATCCACGAAGGTGATTGCTCC |
| qPCR porcine B2M fw | TTCACACCGCTCCAGTAG |
| qPCR porcine B2M rv | CCAGATACATAGCAGTTCAGG |
| qPCR dCas9 reconstitution fw | AGTCTTCACGAGCACATCGC |
| qPCR dCas9 reconstitution rv | CCTTCCCATTACTTTGACGAGTTC |
| (q)PCR MYO7A reconstitution fw | AGAGCAGTGTGAGGCACAAG |
| (q)PCR MYO7A reconstitution rev | CACTGTGGACTCCCCGTCAT |
| (q)PCR MYO7A/Myo7a reconstitution fw | GCCAGAAGAAGAGCAGTGTGA |
| (q)PCR MYO7A/Myo7a reconstitution rv | TGCCGATGATGAAGTGCAGC |
| (q)PCR mMyo7b fw | GGGACACAAGTACAGGAAGGA |
| (q)PCR mMyo7b rv | GCGTTCAAAGCCCACTAGG |
| (q)PCR hMyo7b fw | AAGGACTACGCCCACATCCG |
| (q)PCR hMyo7b rv | CTGAGGCGTCCAGGTTCTC |
| ITR2 fw | GGAACCCCTAGTGATGGAGTT |
| ITR2 rev | CGGCCTCAGTGAGCGA |
| (q)PCR 5'MYO7A fw | ACAAATTCTCAGGCCCCAGT |
| (q)PCR 5'MYO7A rev | TCTCTGGGAAGGACAGCTCC |
| (q)PCR 3'MYO7A fw | GACGCCTTCGTAAAGGGGAT |
| (q)PCR 3'MYO7A rev | CCTGGGAGGGAGGCTTGTAA |
| Myo7a Rx KO deletion (PS1) fw | TTCTCTGCTCGAGGTGAACCC |
| Myo7a Rx KO deletion (PS1) rv | AAAGGCATCTCGCACATCCAG |
| MYO7A Rx KO E14/15 (PS2) fw | GAACTCACAGCACAAGCTCAATG |
| MYO7A Rx KO E14/15 (PS2) rv | TTCTTCTCCAGGAAGCCTTGACTC |
| MYO7A Rx KO E9/12 gDNA (PS3) fw | TACACATGGGCAATCTGCAG |
| MYO7A Rx KO E9/12 gDNA (PS3) rv | AAAGATGTCCAGGAGACCGATG |
| MYO7A Rx KO E9/12 cDNA (PS3) fw | GATCCTGGAAGCATTTGGGA |
| MYO7A Rx KO E9/12 gDNA (PS3) fw | GATGACGTTCATAGGCCGGT |
